# The C-terminal RRM/ACT domain is crucial for fine-tuning the activation of ‘long’ RelA-SpoT Homolog enzymes by ribosomal complexes

**DOI:** 10.1101/849810

**Authors:** Hiraku Takada, Mohammad Roghanian, Victoriia Murina, Ievgen Dzhygyr, Rikinori Murayama, Genki Akanuma, Gemma C. Atkinson, Abel Garcia-Pino, Vasili Hauryliuk

## Abstract

The (p)ppGpp-mediated stringent response is a bacterial stress response implicated in virulence and antibiotic tolerance. Both synthesis and degradation of the (p)ppGpp alarmone nucleotide are mediated by RelA-SpoT Homolog (RSH) enzymes which can be broadly divided in two classes: single-domain ‘short’ and multi-domain ‘long’ RSH. The regulatory ACT (Aspartokinase, Chorismate mutase and TyrA) / RRM (RNA Recognition Motif) domain is a near-universal C-terminal domain of long RSHs. Deletion of RRM in both monofunctional (synthesis-only) RelA as well as bifunctional (i.e. capable of both degrading and synthesising the alarmone) Rel renders the long RSH cytotoxic due to overproduction of (p)ppGpp. To probe the molecular mechanism underlying this effect we characterised *Escherichia coli* RelA and *Bacillus subtilis* Rel RSHs lacking RRM. We demonstrate that, first, the cytotoxicity caused by the removal of RRM is counteracted by secondary mutations that disrupt the interaction of the RSH with the starved ribosomal complex – the ultimate inducer of (p)ppGpp production by RelA and Rel – and, second, that the hydrolytic activity of Rel is not abrogated in the truncated mutant. Therefore, we conclude that the overproduction of (p)ppGpp by RSHs lacking the RRM domain is not explained by a lack of auto-inhibition in the absence of RRM or/and a defect in (p)ppGpp hydrolysis. Instead, we argue that it is driven by misregulation of the RSH activation by the ribosome.

## 1 Introduction

Bacteria employ diverse mechanisms to sense and respond to stress. One such mechanism is the stringent response – a near-universal stress response orchestrated by hyper-phosphorylated derivatives of housekeeping nucleotides GDP and GTP: guanosine tetraphosphate (ppGpp) and guanosine pentaphosphate (pppGpp), collectively referred to as (p)ppGpp (Hauryliuk et al., 2015;Liu et al., 2015;Steinchen and Bange, 2016). Since the stringent response and (p)ppGpp-mediated signalling are implicated in virulence, antibiotic resistance and tolerance (Dalebroux et al., 2010;Dalebroux and Swanson, 2012;Hauryliuk et al., 2015), this stress signalling system has been recently targeted for development of new anti-infective compounds (Kushwaha et al., 2019).

Both synthesis and degradation of (p)ppGpp is mediated by RelA/SpoT Homolog (RSH) enzymes. RSHs can be broadly divided into two classes: ‘long’ multi-domain and ‘short’ single-domain factors (Atkinson et al., 2011;Jimmy et al., 2019). In the majority of bacteria, including model Gram-positive bacterial species *Bacillus subtilis*, the long multi-domain RSHs are represented by one bifunctional enzyme, Rel (Mittenhuber, 2001;Atkinson et al., 2011). Beta- and Gammaproteobacteria, such as *Escherichia coli*, encode two long RSH factors – RelA and SpoT – which are the products of gene duplication and diversification of the ancestral *rel* stringent factor (Mittenhuber, 2001;Atkinson et al., 2011;Hauryliuk et al., 2015). *E. coli* RelA is the most well-studied long RSH. RelA is a dedicated sensor of amino acid starvation with strong (p)ppGpp synthesis activity that is induced by ribosomal complexes harbouring cognate deacylated tRNA in the A-site, so-called ‘starved’ ribosomal complexes (Haseltine and Block, 1973). Unlike RelA, which lacks (p)ppGpp hydrolysis activity (Shyp et al., 2012), Rel and SpoT can both synthesise and degrade (p)ppGpp (Xiao et al., 1991;Avarbock et al., 2000). Similarly to RelA – and to the exclusion of SpoT – (p)ppGpp synthesis by Rel is strongly activated by starved ribosomes (Avarbock et al., 2000). In addition to long RSHs, bacteria often encode single domain RSH enzymes: Small Alarmone Synthetases (SAS) and Small Alarmone Hydrolases (SAH) (Atkinson et al., 2011;Jimmy et al., 2019), such as RelQ and RelP in the Firmicute bacterium *B. subtilis* (Nanamiya et al., 2008).

Long RSHs are universally comprised of two functional regions: the catalytic N-terminal domains (NTD) and the regulatory C-terminal domains (CTD) (**Figure 1A**) (Atkinson et al., 2011). The NTD region comprises the (p)ppGpp hydrolase domain (HD; enzymatically inactive in RelA) and the (p)ppGpp synthetase domain (SYNTH) linked by an α-helical region that regulates the allosteric crosstalk between both domains (Tamman et al., 2019). The CTD encompasses four domains: the Thr-tRNA synthetase, GTPase and SpoT domain (TGS), the Helical domain, the Zing Finger Domain (ZFD) (equivalent to Ribosome-InterSubunit domain, RIS, as per (Loveland et al., 2016), or Conserved Cysteine, CC, as per (Atkinson et al., 2011)), and, finally, the RNA Recognition Motif domain (RRM) (equivalent to Aspartokinase, Chorismate mutase and TyrA, ACT, as per (Atkinson et al., 2011)). When Rel/RelA is bound to a starved ribosomal complex, the TGS domain inspects the deacylated tRNA in the A-site and the TGS domain interacts directly with the 3’ CCA end of the A-site tRNA (Arenz et al., 2016;Brown et al., 2016;Loveland et al., 2016). The conserved histidine 432 residue of *E. coli* RelA mediating this interaction is crucial for activation of RelA’s enzymatic activity by the 3’ CCA (Winther et al., 2018). Both ZFD and RRM interact with the A-site finger (ASF) of the 23S ribosomal RNA (Arenz et al., 2016;Brown et al., 2016;Loveland et al., 2016), and in *E. coli* RelA this contact is crucial for efficient recruitment to and activation by starved ribosomal complexes (Kudrin et al., 2018).

**Figure 1.**
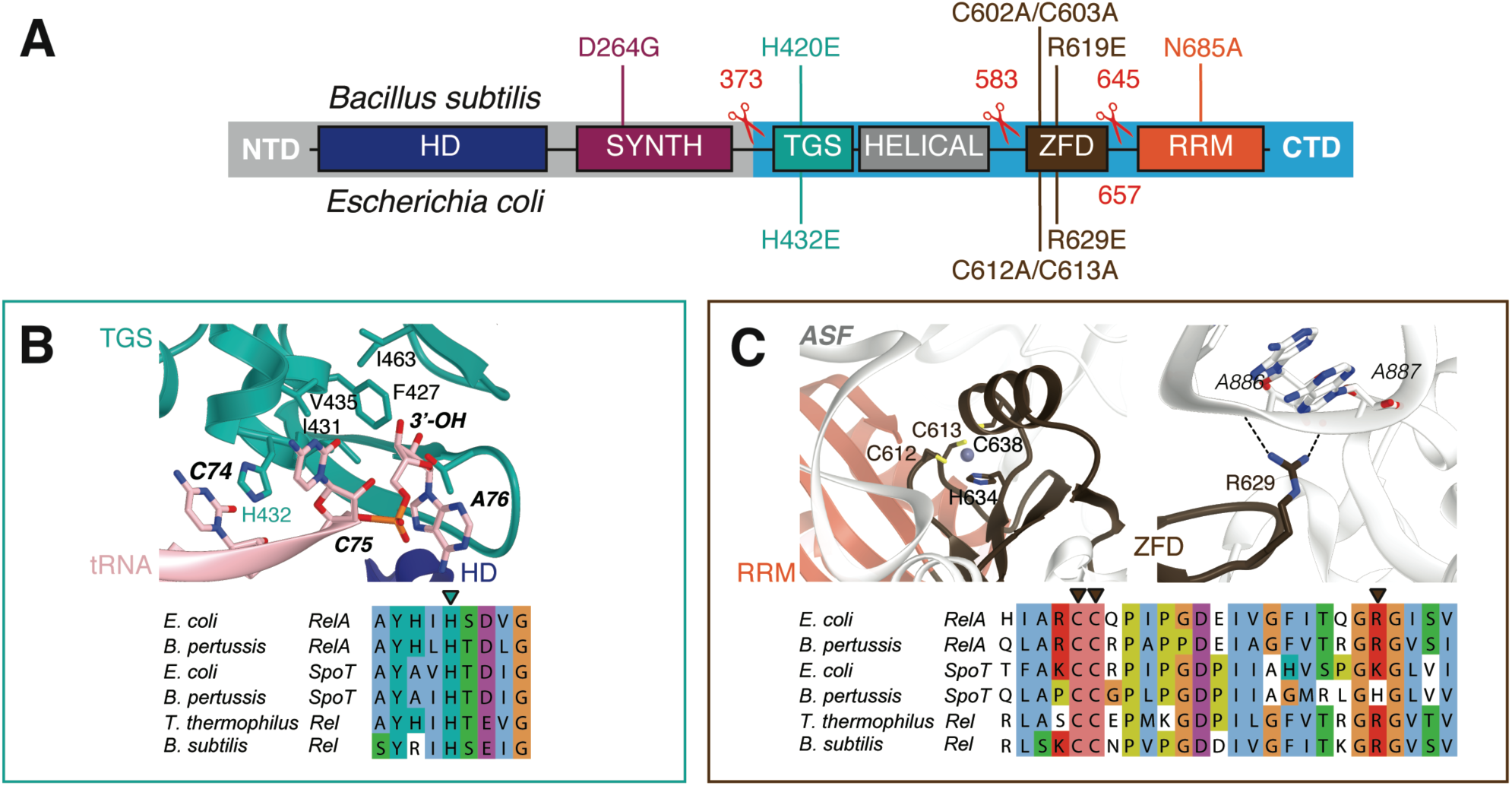
Domain structure of ‘long’ ribosome-associated RSHs Rel and RelA. (**A**) The NTD region contains (p)ppGpp hydrolysis (HD) and (p)ppGpp synthesis (SYNTH) NTD domains. TGS (ThrRS, GTPase and SpoT), Helical, ZFD (Zinc Finger Domain) and RRM (RNA Recognition Motif) domains comprise the regulatory CTD region. Mutations and truncations of *B. subilis* Rel and *E. coli* RelA used in this study are indicated above and below the domain schematics, respectively. (**B**) Conservation and structural environment of mutations in the TGS domain used in the current study. (**C**) Conservation and structural environment of mutations in the RRM domain used in the current study. The 3D structures are as per from Loveland and colleagues (Loveland et al., 2016), RDB accession number 5KPX.

While the NTD is responsible for the enzymatic function of RSHs, the CTD senses the starved ribosomal complex and transmits the signal to activate NTD-mediated (p)ppGpp synthesis by Rel/RelA (Agirrezabala et al., 2013;Arenz et al., 2016;Brown et al., 2016;Loveland et al., 2016). Since removal of the CTD increases the rate of (p)ppGpp production by Rel/RelA in the absence of ribosomes or starved complexes, the CTD was proposed to mediate the auto-inhibition of the NTD synthetase activity, thus precluding uncontrolled production of cytotoxic (p)ppGpp (Schreiber et al., 1991;Gropp et al., 2001;Mechold et al., 2002;Avarbock et al., 2005;Jiang et al., 2007;Yang et al., 2019).

The specific focus of this study is the C-terminal RRM/ACT domain of ribosome-associated RSH RelA and Rel. The RRM is absent in RelA enzymes from *Methylotenera mobilitas, Elusimicrobium minutum*, *Francisella philomiraga* and *Francisella tularensis* (Atkinson et al., 2011). The only experimentally characterised representative amongst these is *F. tularensis* RelA (Wilkinson et al., 2015). In a reconstituted biochemical system, the factor behaves similarly to *E. coli* RelA, i.e. it has very low synthesis activity by itself and is potently activated by the ribosome. Conversely, deletion of the RRM domain in factors that naturally possess it leads to inhibition of growth (Gratani et al., 2018;Ronneau et al., 2019;Turnbull et al., 2019) that is mediated by over-production of (p)ppGpp in the cell, as shown for *Caulobacter crescentus* Rel (Ronneau et al., 2019) and *E. coli* RelA (Turnbull et al., 2019). The exact molecular mechanism of misregulation remains unclear. Deletion of RRM in bifunctional *C. crescentus* Rel leads to compromised hydrolase activity (Ronneau et al., 2019), while overproduction of (p)ppGpp by monofunctional *E. coli* RelA^ΔRRM^ was suggested to be due to upregulated constitutive synthesis activity, conceivably due to defective auto-inhibition of the NTD synthetase domain by the CTD (Turnbull et al., 2019).

In this report, we inspected the possible role of the ribosome in overproduction of (p)ppGpp by ΔRRM variants of long RSHs in the cell. By characterising versions of *E. coli* RelA and *B. subtilis* Rel, we demonstrate that the cytotoxicity of mutant RSH variants is strictly dependent on the interaction with the ribosome and deacylated tRNA, and, therefore, cannot be explained by defects in intra-molecular regulation alone.

## 2 Materials and Methods

### 2.1 Multiple sequence alignment

Sequences were aligned with MAFFT v7.164b with the L-ins-i strategy (Katoh and Standley, 2013), and alignments were visualised with Jalview (Waterhouse et al., 2009).

### 2.2 Construction of bacterial strains and plasmids

The strains and plasmids used in this study are listed in **Supplementary Tables 1-3**. Oligonucleotides used in this study are provided in **Supplementary Table 4**. A detailed description of strain construction is provided in the *Supplementary Material*.

### 2.3 Growth assays

*E. coli* BW25113 cells were transformed with expression constructs either based on a high-copy IPTG inducible vector pUC derivative pMG25 (pMG25::*relA* (Turnbull et al., 2019), pNDM220::*relA^ΔRRM^*, pNDM220::*spoR* or pNDM220::*spoT^ΔRRM^*) or on a low-copy IPTG inducible vector, mini R1 plasmid pNDM220 which is present in one to two copies per chromosome (Molin et al., 1979) (pNDM220::*relA*, pNDM220::*relA^ΔRRM^*, pNDM220::*relA^ΔRRM-H432E^*, pNDM220::*relA^ΔRRM-R629E^* or pNDM220::*relA^ΔRRM-C612A/C613A^*). For solid medium growth assays, ten-fold serial dilutions of overnight LB cultures were spotted onto LB agar supplemented with 30 µg/mL ampicillin and 1 mM IPTG. For liquid medium growth assays, thousand-fold dilutions of the overnight LB cultures were made in liquid LB supplemented with 30 µg/mL ampicillin and 1 mM IPTG, seeded on a 100-well honeycomb plate (Oy Growth Curves AB Ltd, Helsinki, Finland), and plates incubated in a Bioscreen C (Labsystems, Helsinki, Finland) at 37 °C with continuous medium shaking. *B. subtilis* strains were pre-grown on LB plates lacking the IPTG inducer overnight (10 hours) at 30 °C. Fresh individual colonies were used to inoculate filtered LB medium in the presence of indicated concentrations of IPTG and OD_600_ adjusted to 0.01. The cultures were seeded on a 100-well honeycomb plate (Oy Growth Curves AB Ltd, Helsinki, Finland), and plates were incubated in a Bioscreen C (Labsystems, Helsinki, Finland) at 37 °C with continuous medium shaking.

Growth rates (µ_2_) were calculated as slopes of linear regression lines through log_2_-transformed OD_600_ data points.

### 2.4 Preparation of polyclonal anti-Rel antiserum

The entire coding region of the *B. subtilis rel* gene was amplified by PCR using the synthetic oligonucleotide pQErelA_F and pQErelA_R containing a BamHI site and *B. subtilis* genomic DNA as a template. The resulting PCR fragment was cut with BamHI and then inserted into the BamHI sites of pQE60 (Qiagen), yielding plasmid pQErelA. pOErelA was transformed into *E. coli* M15 (pREP4) (Qiagen), fresh transformants were inoculated into LB liquid culture (1000 mL) with 100 µg/mL ampicillin and grown at 37 °C with vigorous shaking. At OD_600_ of 0.8 expression of Rel induced with 1 mM IPTG (final concentration). After 3 hours of expression the cells were harvested by centrifugation, resuspended in buffer A (500 mM NaCl, 50 mM Tris-HCl pH 8.0) supplemented with 2 mM PMSF and lysed by sonication. Rel-His_6_ inclusion bodies were collected by centrifugation, resuspended in buffer A supplemented with 8 M guanidine hydrochloride (= buffer B) and loaded onto an Ni-NTA agarose column (QIAGEN) pre-equilibrated in the same buffer. The column was washed with buffer B supplemented with 10 mM imidazole, and the protein was eluted with a 100-400 mM imidazole gradient in buffer B. The fractions containing Rel-His_6_ was dialyzed against buffer A at 4 °C overnight. Aggregated Rel-His_6_ protein was collected by centrifugation, resuspended in buffer A supplemented with 6 M urea and used for rabbit immunization. Rabbit serum was used as a polyclonal anti-Rel antibody.

### 2.5 Sucrose gradient fractionation and Western blotting

*B. subtilis* strains were pre-grown on LB plates overnight at 30 °C. Fresh individual colonies were used to inoculate 200 mL LB cultures that were grown at 37 °C. At OD_600_ of 0.2 amino acid starvation was induced by addition of isoleucyl tRNA synthetase inhibitor mupirocin (dissolved in DMSO, AppliChem) to final concentration of 700 nM for 20 minutes. As a mock control, a separate culture was treated with the same amount of DMSO. After 20 minutes the cells were collected by centrifugation (8,000 rpm, 5 minutes, JLA-16.25 Beckman Coulter rotor), dissolved in 0.5 mL of HEPES:Polymix buffer (5 mM Mg(OAc)_2_) supplemented with 2 mM PMSF, lysed using FastPrep homogenizer (MP Biomedicals) by four 20 seconds pulses at speed 6.0 mp/sec with chilling on ice for 1 minutes between the cycles), and clarified by ultracentrifugation (14,800 rpm for 20 minutes, Microfuge 22R centrifuge Beckman Coulter, F241.5P rotor). Clarified cell lysates were loaded onto 10-35% sucrose gradients in HEPES:Polymix buffer pH 7.5 (5 mM Mg^2+^ final concentration), subjected to centrifugation (36,000 rpm for 3 hours at 4 °C, SW-41Ti Beckman Coulter rotor) and analysed using Biocomp Gradient Station (BioComp Instruments) with A_260_ as a readout.

For Western blotting 0.5 mL fractions were supplemented with 1.5 mL of 99.5% ethanol, precipitated overnight at –20 °C. After centrifugation at 14,800 rpm for 30 minutes at 4 °C the supernatants were discarded and the samples were dried. The pellets were resuspended in 40 µL of 2xSDS loading buffer (100 mM Tris-HCl pH 6.8, 4% SDS (w/v) 0.02% Bromophenol blue, 20% glycerol (w/v) 4% β-mercaptoethanol), resolved on the 8% SDS PAGE and transferred to nitrocellulose membrane (Trans-Blot Turbo Midi Nitrocellulose Transfer Pack, Bio-Rad, 0.2 µm pore size) with the use of a Trans-Blot Turbo Transfer Starter System (Bio-Rad) (10 minutes, 2.5A, 25V). Membrane blocking was done for one hour in PBS-T (1xPBS 0.05% Tween-20) with 5% w/v nonfat dry milk at room temperature. Rel was detected using anti-Rel primary combined with goat anti-rabbit IgG-HRP secondary antibodies. All antibodies were used at 1:10,000 dilution. ECL detection was performed using WesternBright^TM^ Quantum (K-12042-D10, Advansta) Western blotting substrate and an ImageQuant LAS 4000 (GE Healthcare) imaging system.

### 2.6 Expression and purification of *E. coli* RelA and *B. subtilis* Rel

Wild type and H432E mutant variants of *E. coli* RelA were expressed and purified as described earlier (Turnbull et al., 2019).

Wild type and mutant variants of *B. subtilis* Rel were overexpressed in freshly transformed *E. coli* BL21 DE3 Rosetta (Novagen). Fresh transformants were inoculated to final OD_600_ of 0.05 in the LB medium (800 mL) supplemented with 100 µg/mL kanamycin. The cultures were grown at 37 °C until an OD_600_ of 0.5, induced with 1 mM IPTG (final concentration) and grown for additional 1.5 hour at 30 °C. The cells were harvested by centrifugation and resuspended in buffer A (750 mM KCl, 5 mM MgCl_2_, 40 µM MnCl_2_, 40 µM Zn(OAc)_2_, 1 mM mellitic acid (Tokyo Kasei Kogyo Co., Ltd.), 20 mM imidazole, 10% glycerol, 4 mM β-mercaptoethanol, 25 mM HEPES:KOH pH 8) supplemented with 0.1 mM PMSF and 1 U/mL of DNase I. Cells were lysed by one passage through a high-pressure cell disrupter (Stansted Fluid Power, 150 MPa), cell debris was removed by centrifugation (25,000 rpm for 40 min, JA-25.50 Beckman Coulter rotor) and clarified lysate was taken for protein purification.

To prevent possible substitution of Zn^2+^ ions in Rel’s Zn-finger domain for Ni^2+^ during purification on an Ni-NTA metal affinity chromatography column (Block et al., 2009), a 5 mL HisTrap HP column was stripped from Ni^2+^ in accordance with manufacturer’s recommendations, washed with 5 column volumes (CV) of 100 mM Zn(OAc)_2_ pH 5.0 followed by 5 CV of deionized water. Clarified cell lysate was filtered through a 0.2 µm syringe filter and loaded onto the Zn^2+^-charged HisTrap 5 mL HP column pre-equilibrated in buffer A. The column was washed with 5 CV of buffer A, and the protein was eluted with a linear gradient (6 CV, 0-100% buffer B) of buffer B (750 mM KCl, 5 mM MgCl_2_, 40 µM MnCl_2_, 40 µM Zn(OAc)_2_, 1 mM mellitic acid, 500 mM imidazole, 10% glycerol, 4 mM β-mercaptoethanol, 25 mM HEPES:KOH pH 8). Mellitic acid forms highly ordered molecular networks when dissolved in water (Inabe, 2005) and it was shown to promote the stability of *Thermus thermophilus* Rel (Van Nerom et al., 2019). Fractions most enriched in Rel (≈25-50% buffer B) were pooled, totalling approximately 5 mL. The sample was loaded on a HiLoad 16/600 Superdex 200 pg column pre-equilibrated with a high salt buffer (buffer C; 2 M NaCl, 5 mM MgCl_2_, 10% glycerol, 4 mM β-mercaptoethanol, 25 mM HEPES:KOH pH 8). The fractions containing Rel were pooled and applied on HiPrep 10/26 desalting column (GE Healthcare) pre-equilibrated with storage buffer (buffer D; 720 mM KCl, 5 mM MgCl_2_, 50 mM arginine, 50 mM glutamic acid, 10% glycerol, 4 mM β-mercaptoethanol, 25 mM HEPES:KOH pH 8). Arginine and glutamic acid were added to improve protein solubility and long-term stability (Golovanov et al., 2004). The fractions containing Rel were collected and concentrated in an Amicon Ultra (Millipore) centrifugal filter device (cut-off 50 kDa). To cleave off the His_10_-SUMO tag, 35 µg of His_6_-Ulp1 per 1 mg of Rel were added and the reaction mixture was incubated at room temperature for 15 min. After the His_10_-SUMO tag was cleaved off, the protein was passed though 5 mL Zn^2+^-charged HisTrap HP pre-equilibrated with buffer D. Fractions containing Rel in the flow-through were collected and concentrated on Amicon Ultra (Millipore) centrifugal filter device with 50 kDa cut-off. The purity of protein preparations was assessed by SDS-PAGE and spectrophotometrically (OD_260_/OD_280_ ratio below 0.8 corresponding to less than 5% RNA contamination (Layne, 1957)). Protein preparations were aliquoted, frozen in liquid nitrogen and stored at –80 °C. Individual single-use aliquots were discarded after the experiment.

### 2.7 Negative staining electron microscopy

3.5 µL of 2 µM Rel protein was loaded onto a glow-discharged Cu_300_ grid (TAAB Laboratories Equipment Ltd.) with manually layered 2.9 nm carbon. The sample was incubated on the grid for 1-3 minutes, blotted with Watman filter paper, than twice washed with water and blotted, stained with 1.5% uranyl acetate pH 4.2 for 30 seconds before the final blotting. Grids were dried on the bench and imaged by Talos L 120C (FEI) microscope with 92,000X magnification.

### 2.8 Preparation of 10X Polymix buffer base

The 10X Polymix base was prepared as per (Antoun et al., 2004), with minor modifications. For preparation of the putrescine solution, 100 g of putrescine (1,4-diaminobutane) was dissolved in 600 mL of ddH_2_O at 90 °C, and the pH adjusted with acetic acid to 8.0 (approximately 100 mL of 100% acetic acid). After cooling to room temperature, the pH was adjusted further to 7.6 and the volume was adjusted to the final of 2 L by addition of 1.134 L of ddH_2_O. One 100 mL cup of activated charcoal was added and the slurry was stirred under the hood for 30 minutes. The slurry was filtered through, first, Whatman paper and then through a 0.45 µm BA85 membrane. The final solution was stored at 4 °C in a bottle wrapped in foil since putrescine is photosensitive. The preparation of 2 L of 10X Polymix buffer base used 141.66 g KCl, 5.35 g NH_4_Cl, 21.44 g Mg(OAc)_2_•4H_2_O, 1.47 g CaCl_2_•2H_2_O, 5.092 g spermidine, and 160 ml of putrescine solution (described above). The salts were dissolved in ddH_2_O (≈1,500 mL), then the putrescine solution was added and mixed well. Spermidine was dissolved in a small volume of ddH_2_O and added to the mixture. The pH was adjusted to 7.5 with concentrated acetic acid or 5 M KOH, and after that the volume was adjusted by adding ddH_2_O to 2 L. The buffer was filtered through 0.2 µm nitrocellulose filter (2-3 filters are needed). The resulting 10X Polymix buffer base was aliquoted and stored at –20 °C. The final working HEPES:Polymix buffer was made using the 10X Polymix buffer base, 1M DDT and 1 M HEPES:KOH pH 7.5 and contains 20 mM HEPES:KOH pH 7.5, 2 mM DTT, 5 mM MgOAc_2_, 95 mM KCl, 5 mM NH_4_Cl, 0.5 mM CaCl_2_, 8 mM putrescine, 1 mM spermidine.

### 2.9 Purification of *B. subtilis* 70S ribosomes

*B. subtilis* strain RIK2508 (*trpC2 Δhpf*) strain (Akanuma et al., 2016;Brodiazhenko et al., 2018) was pre-grown on LB plates overnight at 30 °C. Fresh individual colonies were used to inoculate LB liquid cultures (25 × 400 mL) to OD_600_ of 0.05 and grown at 37 °C with vigorous shaking. At OD_600_ 1.2 the cells were pelleted at 4 °C (TLA10.500 (Beckman), 15 min at 5,000-8,000 rcf), resuspended with ice-cold PBS buffer, pelleted again in 50 mL Falcon tubes, frozen with liquid nitrogen and stored at –80 °C. Approximately 20 g of frozen *B. subtilis* cells were resuspended in 50 mL of cell opening buffer (100 mM NH_4_Cl, 15 mM Mg(OAc)_2_, 0.5 mM EDTA, 3 mM β-mercaptoethanol, 20 mM Tris:HCl pH 7.5) supplemented with 1 mU Turbo DNase (Thermo Fisher Scientific), 0.1 mM PMSF and 35 µg/mL lysozyme, incubated on ice for one hour, and opened by three passages on a high-pressure cell disrupter (Stansted Fluid Power) at 220 MPa. Lysed cells were clarified by centrifugation for 40 min at 40,000 rpm (Ti 45 rotor, Beckman), NH_4_Cl concentration was adjusted to 400 mM, and the mixture was filtered through 0.45 µm syringe filters. The filtrated lysate was loaded onto a pre-equilibrated 80 mL CIMmultus QA-80 column (BIA Separations, quaternary amine advanced composite column) at a flow rate of 20 mL/min, and the column washed with 5 CV (CV = 80 mL) of low salt buffer (400 mM NH_4_Cl, 15 mM Mg(OAc)_2_, 3 mM β-mercaptoethanol, 20 mM Tris:HCl pH 7.5). Ribosomes were then eluted in 45 mL fractions by a step gradient to 77% high salt buffer (900 mM NH_4_Cl, 15 mM Mg(OAc)_2_, 3 mM β-mercaptoethanol, 20 mM Tris:HCl pH 7.5) for 5 CV, followed by 100% high salt buffer for 1 CV. The fractions containing ribosomes were pooled, the concentration of NH_4_Cl was adjusted to 100 mM, and the ribosomes were treated with puromycin added to a final concentration of 10 µM. The resultant crude 70S preparation was resolved on a 10-40% sucrose gradient in overlay buffer (60 mM NH_4_Cl, 15 mM Mg(OAc)_2_, 0.25 mM EDTA, 3 mM β-mercaptoethanol, 20 mM Tris:HCl pH 7.5) in a zonal rotor (Ti 15, Beckman, 17 hours at 21,000 rpm). The peak containing pure 70S ribosomes was pelleted by centrifugation (20 hours at 35,000 rpm), and the final ribosomal preparation was dissolved in HEPES:Polymix buffer (20 mM HEPES:KOH pH 7.5, 2 mM DTT, 5 mM Mg(OAc)_2_, 95 mM KCl, 5 mM NH_4_Cl, 0.5 mM CaCl_2_, 8 mM putrescine, 1 mM spermidine (Antoun et al., 2004)). 70S concentration was measured spectrophotometrically (1 OD_260_ corresponds to 23 nM of 70S) and ribosomes were aliquoted (50-100 µL per aliquot), snap-frozen in liquid nitrogen and stored at –80 °C.

### 2.10 Preparation of 70S initiation complexes (70S IC)

Initiation complexes were prepared by as per (Murina et al., 2018), with minor modifications. The reaction mix containing *B. subtilis* 70S ribosomes (final concentration of 6 µM) with *E. coli* IF1 (4 µM), IF2 (5 µM), IF3 (4 µM), ^3^H-fMet-tRNA_i_^fMet^ (8 µM), mRNA MVFStop (8 µM, 5’-GGC**AAGGAGGA**GAUAAGA<u>AUGGUUUUCUAA</u>UA-3’; Shine-Dalgarno sequence is highlighted in bold, ORF is underlined), 1 mM GTP and 2 mM DTT in 1×HEPES:Polymix buffer (20 mM HEPES:KOH pH 7.5, 95 mM KCl, 5 mM NH_4_Cl, 5 mM Mg(OAc)_2_, 0.5 mM CaCl_2_, 8 mM putrescine, 1 mM spermidine, 1 mM DTT (Antoun et al., 2004)) was incubated at 37 °C for 30 min. Then the ribosomes were pelleted through a sucrose cushion (1.1 M sucrose in HEPES:Polymix buffer with 15 mM Mg^2+^) at 50,000 rpm for two hours (TLS-55, Beckman), the pellet was dissolved in HEPES:Polymix buffer (5 mM Mg(OAc)_2_), aliquoted, frozen in liquid nitrogen and stored at –80 °C.

### 2.11 Preparation of ^3^H-labelled pppGpp

3 µM *E. fecalis* RelQ (Beljantseva et al., 2017) were incubated in reaction buffer (18 mM MgCl_2_, 20 mM DTT, 20 M Tris-HCl pH 8.0) together with 8 mM ATP and 5 mM ^3^H-GTP (SA: 100 cpm/pmol) for 2 hours at 37 °C to produce ^3^H-pppGpp. The resultant mixture was loaded on strong anion-exchange column (MonoQ 5/50 GL; GE Healthcare), and nucleotides were resolved by a 0.5-1,000 mM LiCl gradient. Peak fractions containing ^3^H-pppGpp were pooled and precipitated by addition of lithium chloride to a final concentration of 1 M followed by addition of four volumes of ethanol. The suspension was incubated at –80 °C overnight and centrifuged (14,800 rpm, 30 min, 4 °C). The resulting pellets were washed with absolute ethanol, dried, dissolved in 20 mM HEPES-KOH buffer (pH 7.5) and stored at –80 °C.

### 2.12 ^3^H-pppGpp hydrolysis assay

The reaction mixtures contained 140-250 nM Rel, 300 µM ^3^H-pppGpp, 1 mM MnCl_2_, an essential cofactor for Rel’s hydrolysis activity (Avarbock et al., 2000;Mechold et al., 2002;Tamman et al., 2019), all in HEPES:Polymix buffer (5 mM Mg^2+^ final concentration). After preincubation at 37 °C for 3 minutes, the reaction was started by the addition of prewarmed Rel and 5 µL aliquots were taken throughout the time course of the reaction and quenched with 4 µL 70% formic acid supplemented with a cold nucleotide standard (4 mM GTP) for UV-shadowing.

### 2.13 ^3^H-pppGpp synthesis assay

Assays with *E. coli* RelA were performed as described earlier (Kudrin et al., 2018). In the case of *B. subtilis* Rel, the reaction mixtures typically contained 500 nM *B. subtilis* 70S IC(MVF), 140 nM Rel, guanosine nucleoside substrate (300 µM ^3^H-GTP, PerkinElmer), 100 µM pppGpp, 2 µM *E. coli* tRNA^Val^ (Chemical Block), all in HEPES:Polymix buffer (5 mM Mg^2+^ final concentration). After preincubation at 37 °C for 3 minutes, the reaction was started by the addition of prewarmed ATP to the final concentration of 1 mM, and 5 µL aliquots were taken throughout the time course of the reaction and quenched with 4 µL 70% formic acid supplemented with a cold nucleotide standard (4 mM GTP) for UV-shadowing. Individual quenched timepoints were spotted PEI-TLC plates (Macherey-Nagel) and nucleotides were resolved in 1.5 KH_2_PO_4_ pH 3.5 buffer. The TLC plates were dried, cut into sections as guided by UV-shadowing, and ^3^H radioactivity was quantified by scintillation counting in EcoLite™ Liquid Scintillation Cocktail (MP Biomedicals).

## 3 Results

### 3.1 Toxicity of *E. coli* ΔRRM RelA is countered by mutations compromising the interaction with starved ribosomes

As we have shown earlier, low-level ectopic expression of RelA^ΔRRM^ from a low copy number pNDM220 plasmid under the control of a P_A1/O4/O3_ promoter has a more pronounced inhibitory effect on *E. coli* growth in comparison with expression of the full-length protein (Turnbull et al., 2019). We tested whether P_A1/O4/O3_-driven high-level expression of RelA^ΔRRM^ from a high-copy pUC derivative pMG25 would cause a more pronounced growth defect (**Figure 2A**). For comparison, we tested the effects of expression of the second *E. coli* RSH enzyme – SpoT – using both the full-length and the ΔRRM variants. Even in the absence of the IPTG inducer, leaky expression of RelA^ΔRRM^ has a dramatic effect on *E. coli* growth, while the full-length protein does not have an effect. In the presence of 50 µM IPTG, both full-length and RelA^ΔRRM^ inhibit the growth, although the latter has a stronger effect; induction with 500 µM IPTG completely abrogates the growth in both cases. While expression of SpoT has a detectable inhibitory effect at 500 µM IPTG, the effect is the same for full-length and SpoT^ΔRRM^. Since even leaky expression of RelA^ΔRRM^ inhibits growth, we concluded that this high-level expression system is ill-suited for follow-up microbiological investigations. Therefore, to test whether the toxicity of RelA^ΔRRM^ in *E. coli* is dependent on the interaction with starved complexes, we used the pNDM220-based low-level expression system used previously (Turnbull et al., 2019). Guided by the recent cryo-EM reconstructions of RelA (Arenz et al., 2016;Brown et al., 2016;Loveland et al., 2016), we designed a set of mutations that will specifically disrupt RelA’s interaction with starved ribosomal complexes.

**Figure 2.**
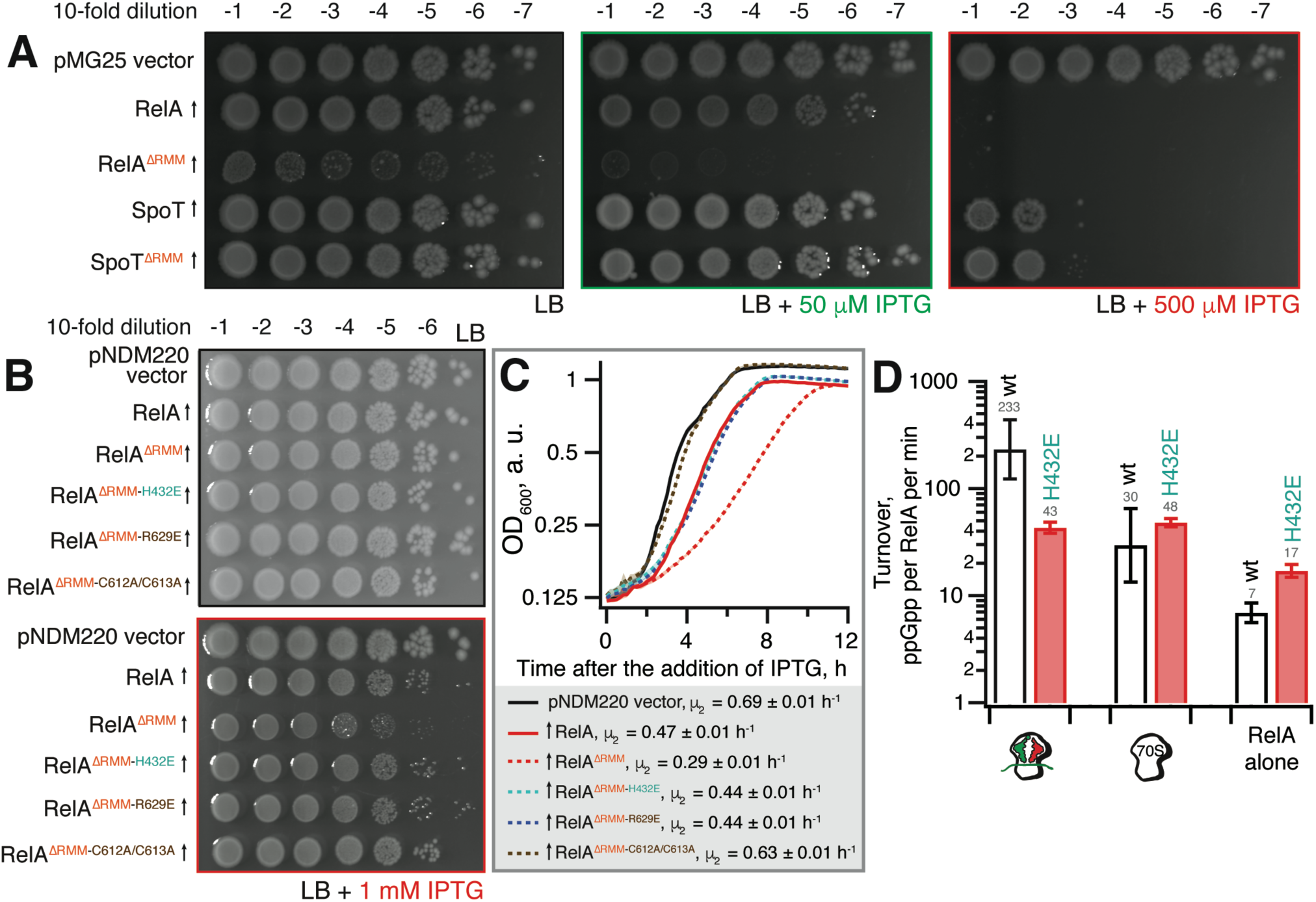
The toxicity of ΔRRM *E. coli* RelA is mitigated by mutations compromising interactions with tRNA and the ribosome. (**A**) Wild-type *E. coli* BW25113 cells were transformed with either the empty high-copy IPTG-inducible pMG25 plasmid vector or pMG25-based constructs expressing wild-type and ΔRRM versions of *E. coli* RelA and SpoT. Up-pointing arrows indicate induction of expression. Ten-fold serial dilutions of overnight LB cultures were made and spotted onto LB agar supplemented with 100 µg/mL ampicillin and either 0, 50 or 500 µM IPTG. The plates were incubated at 37 °C and scored after 18 hours. (**B**) *E. coli* BW25113 cells were transformed either with the empty low-copy IPTG-inducible pNDM220 vector or pNDM220-based constructs expressing wild-type and mutant versions of *E. coli* RelA as indicated on the figure. Ten-fold serial dilutions of overnight LB cultures were made and spotted onto LB agar supplemented with 30 µg/mL ampicillin and 1 mM IPTG. As a plating control the same dilutions of the overnight cultures were spotted on LB agar supplemented with 30 µg/mL ampicillin but no IPTG. The plates were incubated at 37 °C and scored after 18 hours. (**C**) Thousand-fold dilutions of the same overnights were made in LB supplemented with 30 µg/mL ampicillin and 1 mM IPTG, and growth at 37 °C was monitored using the Bioscreen C growth curve analysis system. The growth rates (µ^2^) were calculated from three independent measurements and the error bars represent standard errors. (**D**) H432E TGS *E. coli* RelA is not activated by deacylated tRNA on the ribosome. The synthetase activity of 30 nM wild type and H432E *E. coli* RelA was assayed in the presence of 1 mM ATP, 300 µM ^3^H GDP and 100 µM ppGpp in HEPES:Polymix buffer, pH 7.5, 37 °C, 5 mM Mg^2+^. As indicated on the figure, the reaction mixture was supplemented either with 2 µM vacant 70S ribosomes or with an *in situ* assembled starved ribosomal complex (2 µM vacant 70S combined with 2 µM mRNA(MV), 2 µM *E. coli* tRNA_i_^fMet^ and 2 µM *E. coli* tRNA^Val^). The error bars represent standard deviations of the turnover estimates by linear regression using four data points.

To disrupt the interaction between RelA and the tRNA, we adopted the H432E mutation in the TGS domain that was earlier shown to specifically abrogate the recognition of the 3’ CCA end of the A/R tRNA (Winther et al., 2018). This conserved histidine residue stacks between the two cytosine bases and hydrogen-bonds the phosphodiester backbone (**Figure 1B**). Replacing it with glutamic acid introduces a charge repulsion effect as well as a steric clash. To disrupt the interaction with the ribosome we used mutations in the ZFD: R629E as well as a double substitution C602A C603A; both mutants are expected to compromise the recognition of the 23S rRNA ASF element that is crucial for RelA recruitment (Kudrin et al., 2018). The conserved double motif docks the ZFD α-helix into the major groove of the ASF; replacement by alanine is expected to abrogate this interaction (**Figure 1C**). The conserved arginine 629 residue is in close proximity to A886-A887 and A885-A886 phosphodiester bonds (3.5Å and 5Å, respectively), and, therefore, the R629E substitution is expected to cause electrostatic repulsion.

Low-level expression of full-length RelA has a minor, but detectable growth inhibitory effect both when tested on solid LB agar media (**Figure 2B**) and in liquid LB cultures (growth rate, µ_2_, decreases from 0.69 (vector) to 0.47 h^-1^) (**Figure 2C**). Importantly, the spotting control on solid LB media lacking IPTG shows that the size of the inoculum is not affected by potential leaky expression in the overnight culture (**Figure 2B**). Deletion of the RRM renders RelA significantly more toxic (growth rate decreases to 0.29 h^-1^), in good agreement with the accumulation of (p)ppGpp upon expression of the construct (Turnbull et al., 2019). The effect is countered by TGS H432E, ZFD R629E and even more so by the C612A C613A substitutions (**Figure 2BC**). RelA^ΔRRM^ expression is equally toxic in the Δ*relA* background and the effect is similarly countered by H432E, R629E and C612A C613A substitutions (**Supplementary Figure S1**), demonstrating that the growth inhibition is independent of the functionality of the endogenous RelA stringent factor. Finally, we directly confirmed the lack of activation by deacylated tRNA in the case of H432E *E. coli* RelA using biochemical assays (**Figure 2D**).

Taken together, our results suggest that RelA^ΔRRM^ toxicity in *E. coli* is dependent on the functionality of the interaction with starved ribosomes. To test the generality of this hypothesis, we next characterised *B. subtilis* Rel lacking the RRM domain.

### 3.2 Toxicity of *B. subtilis* ΔRRM Rel expressed in the ppGpp^0^ background is mediated by (p)ppGpp synthesis and is countered by mutations compromising the interaction with starved ribosomes

We expressed ΔRRM Rel under the control of an IPTG-inducible P*_hy-spank_* promotor (Britton et al., 2002) in ppGpp^0^ (Δ*rel* Δ*relP* Δ*relQ*) (Nanamiya et al., 2008) or Δ*rel B. subtilis* strains (**Figure 3A**). In the ppGpp^0^ background, inhibition of *B. subtilis* growth on LB plates is a marker of toxic (p)ppGpp overproduction. In the Δ*rel* background, (p)ppGpp is overproduced by a SAS in the absence of Rel’s hydrolytic activity, causing a growth defect in *B. subtilis* (Nanamiya et al., 2008) and *S. aureus* (Geiger et al., 2014). Therefore, this experiment tests the complementation of hydrolase function of Rel that manifests in improved growth.

**Figure 3.**
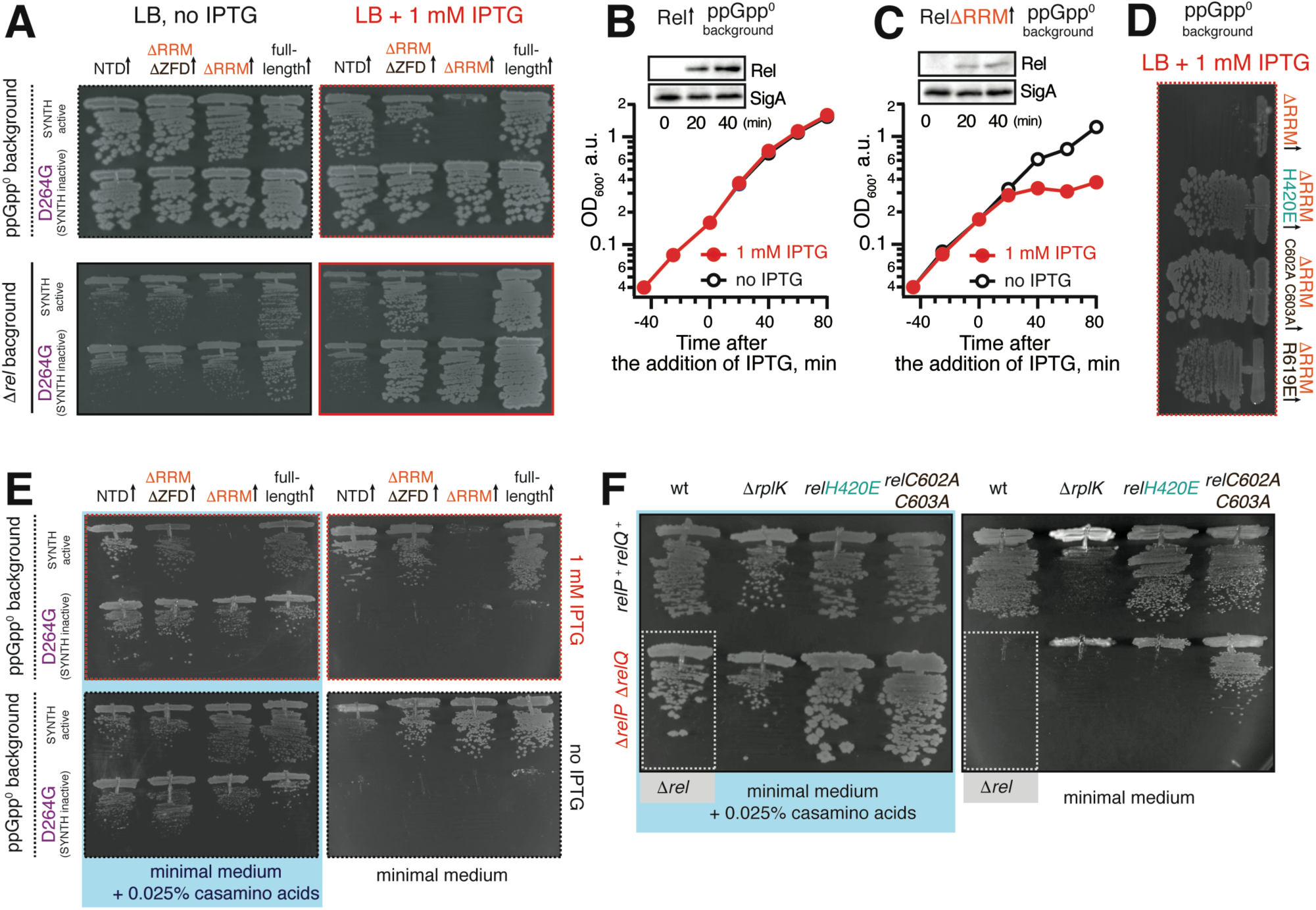
Deletion of the regulatory RRM domain leads to *B. subtilis* Rel toxicity due to ribosome-dependent (p)ppGpp overproduction. (**A**) Full-length (VHB155 and VHB183) as well as C-terminally truncated Rel variants [synthesis-competent ΔRRM (VHB159 and VHB184), ΔRRMΔZFD (VHB160 and VHB185) and Rel^NTD^ (VHB161 and VHB186), and the corresponding synthesis-inactive D264G mutants VHB162-164; VHB187-190] were expressed in either ppGpp^0^ (upper row; test for toxicity mediated by (p)ppGpp accumulation) or Δ*rel* (lower row; test for HD functionality) *B. subtilis* growing on solid LB medium. Up-pointing arrows (↑) indicate ectopic expression. (**B**-**C**) Expression of Rel^ΔRRM^ causes a growth defect in liquid culture. Either wild-type *rel* (VHB183) (**B**) or *rel^ΔRRM^* mutant (VHB184) (**C**) were expressed in ppGpp^0^ background grown in liquid LB medium at 37 °C. Protein expression was induced by IPTG added to final concentration of 1 mM to exponentially growing bacterial cultures at OD_600_ 0.2. Protein expression was monitored by Western blotting using anti-Rel antibodies (see also **Supplementary Figure S2H**). (**D**) The toxicity of mutant versions of Rel^ΔRRM^ tested in ppGpp^0^ *B. subtilis* growing on solid LB medium: wild type Rel^ΔRRM^ (VHB184), H420E (VHB231) defective in recognition of the tRNA 3’ CCA end, and ZFD mutants C602A C603A (VHB233) and R619E (VHB281) defective in 70S binding. LB plates were scored after 18 hour incubation at 37 °C. (**E**) C-terminally truncated Rel variants (either synthesis-competent (VHB155, VHB159-161) or synthesis-inactive D264G mutant versions (VHB156, VHB162-164)) were expressed in ppGpp^0^ *B. subtilis* growing on either solid minimal medium or solid minimal medium supplemented with 0.025% casamino acids. Plates were scored after 36 hours incubation at 37 °C. Importantly, prior to experiment all strains were pre-cultured on solid minimal medium supplemented with 0.025% casamino acids. This was done in order to avoid the effects caused by the decreased fitness of the inoculum. (**F**) Synthesis activity of Rel mutants probed by amino acid auxotrophy assays. *B. subtilis* strains were constructed using either *relP^+^ relQ^+^* wild-type 168 (upper row) or *ΔrelP ΔrelP* (lower row) background. The strains either expressed the indicated *rel* mutants (H420E (VHB68) and C602A C603A (VHB148), upper row, and H420E (VHB60), C602A C603A (VHB62), lower row) or contained an additional *ΔrplK* gene disruption (VHB47, *relP^+^ relQ^+^* and VHB49, *ΔrelP ΔrelP*). The ppGpp^0^ mutant strain (*ΔrelP ΔrelP Δrel*, VHB63) was used as a control (highlighted with grey box). The strains were grown on either solid Spizizen minimal medium (right panel) or solid minimal medium supplemented with 0.025% casamino acids (left panel).

Unlike the full-length Rel, the Rel^ΔRRM^ truncation is toxic in the ppGpp^0^ and Δ*rel* backgrounds, both when the growth is followed on plates (**Figure 3A**) and in liquid culture (**Figure 3BC**). To probe the role of the interactions with starved ribosomes in Rel^ΔRRM^ toxicity, we used a set of substitutions in *subtilis* Rel corresponding to those used to study *E. coli* RelA (**Figures 1** and **2**). The toxicity of the Rel^ΔRRM^ mutant is efficiently countered by the H420E substitution in the TGS as well as the R619E and C602A C603A substitutions in the ZFD (**Figure 3D** and **Supplementary Figure S2D**-**F, H**). This strongly suggests that the intact interaction with tRNA and starved ribosomes is essential for the toxicity of Rel^ΔRRM^. For comparison, we tested, full-length Rel, Rel^ΔRRMΔZDF^ C-terminal truncation, as well as the NTD domain region alone. The Rel^ΔRRMΔZDF^ mutant is only slightly toxic in the ppGpp^0^ background and its expression promotes growth in the Δ*rel* background (**Figure 3A**). NTD-alone construct displays no toxicity in the ppGpp^0^ background and does not promote the growth in the Δ*rel* background, suggesting weak – or absent – synthetase activity.

To separate the effects of (p)ppGpp production from the effects of (p)ppGpp degradation, we tested synthesis-deficient SYNTH D264G mutants (Nanamiya et al., 2008) of C-terminally truncated Rel variants (**Figure 3A**). The toxicity of the ΔRRM variant is abolished by the D264G mutation demonstrating that it is, indeed, mediated by (p)ppGpp production and not, for example, through inhibition of protein synthesis via competitive binding to ribosomal A-site (the latter non-enzymatic mechanism of toxicity was shown for *E. coli* RelA^CTD^ (Turnbull et al., 2019)). Both ΔRRM D264G and ΔZFDΔRRM D264G variants promote growth in the Δ*rel* strain suggesting that neither deletion of ΔRRM alone – or both ΔZFD and ΔRRM – abrogates the hydrolysis activity of *B. subtilis* Rel (**Figure 3A**, bottom panel). At the same time expression of the synthesis-inactive D264G Rel^NTD^ has no effect, suggesting that the NTD does not efficiently hydrolyse (p)ppGpp. To test if further truncations of the NTD-only Rel (Rel^1-373^) would induce hydrolytic activity, we tested several additional constructs of *B. subtilis* Rel – Rel^1-336^, Rel^1-196^ and Rel^155^ – but neither of them could rescue the growth effect of *Δrel B. subtilis* (**Supplementary Figure S3**). Finally, the synthesis deficiency of D264G Rel and the lack of H420E Rel activation by deacylated tRNA was confirmed using biochemical assays (**Supplementary Figure S4**).

Taken together, our results demonstrate that i) Rel^ΔRRM^ is toxic in ppGpp^0^ *B. subtilis,* ii) this toxicity requires intact (p)ppGpp synthesis activity of the enzyme iii) it is abrogated by mutations disrupting the interaction with tRNA and starved ribosomes and iv) deletion of the RRM domain does not abrogate the hydrolysis activity of *B. subtilis* Rel.

### 3.3 The synthetase activity of Rel^ΔRRM^, Rel^ΔRRMΔZFD^ and Rel^NTD^ but not the Rel^H420E^ TGS mutant can suppress amino acid auxotrophy of ppGpp^0^ B. subtilis

To test the low-level, non-toxic, synthesis activity of Rel mutants and to validate the effects of point mutations disrupting the interaction of Rel with starved ribosomal complexes, we took advantage of the amino acid auxotrophy phenotype of the ppGpp^0^ *B. subtilis* (Δ*rel* Δ*relP* Δ*relQ*) (Nanamiya et al., 2008)).

When ppGpp^0^ *B. subtilis* is grown on Spizizen minimum medium (Spizizen, 1958) in the absence of casamino acids, neither of the D264G Rel mutants – either full-length or C-terminal truncations – promote growth, both whether or not expression is induced by 1 mM IPTG (**Figure 3E**, right panels). Full induction of NTD expression with 1 mM IPTG near-completely supressed the auxotrophy phenotype (**Figure 3E**, top right panel), while the leaky expression in the absence of IPTG results in weak, but detectable suppression (**Figure 3E**, bottom right panel). This demonstrates that *B. subtilis* NTD has a weak net-synthesis activity. The Rel^ΔRRM^ is, as expected, highly toxic when expression is induced by IPTG; conversely, low-level leakage expression efficiently supresses the amino acid auxotrophy phenotype. Removal of both RRM and ZFD domains renders the protein non-toxic. It is not trivial to reconcile this effect with the idea that removal of the RRM renders the protein toxic due to lack of auto-inhibition: one would expect that the additional removal of the ZFD domain would further compromise the CTD-mediated negative control in Rel^ΔRRM^.

We next used the auxotrophy assay to test the effects of the H420E TGS and C602A C603A ZFD substitutions on the activity of Rel expressed from the native genomic locus under the control of the native promotor (**Figure 3F**). As a positive control we used a strain lacking the genomic copy of *rplK* (*relC*) encoding ribosomal protein L11. This ribosomal element is essential for *E. coli* RelA activation by starved ribosomal complexes (Parker et al., 1976;Wendrich et al., 2002;Shyp et al., 2012) as well as for cellular functionality of *C. crescentus* Rel (Boutte and Crosson, 2011). The ppGpp^0^ strain expressing H420E Rel fails to grow on the minimum media, reinforcing the crucial role of that this residue, while the C602A C603A can sustain the growth, suggesting that this substitution does not completely abrogate the activity.

### 3.4 RRM deletion and ZFD mutations destabilise B. subtilis Rel binding to starved ribosomal complexes

Next, we probed the ribosomal association of ΔRRM and full-length Rel expressed in the ppGpp^0^ background using a centrifugation sucrose gradient followed by Western blotting using antiserum against native, untagged *B. subtilis* Rel (**Figure 4A**). Since deacylated tRNA promotes ribosomal recruitment of *E. coli* RelA (Agirrezabala et al., 2013;Kudrin et al., 2017), we probed the association of wild type and mutant Rel variants with the ribosome both under exponential growth and upon acute isoleucine starvation induced by the isoleucyl tRNA synthetase inhibitor antibiotic mupirocin (pseudomonic acid) (Thomas et al., 2010). In good agreement with the cryo-EM structures detailing multiple contacts between the RRM and the starved complex and therefore suggesting an importance of this element in ribosomal recruitment (Arenz et al., 2016;Brown et al., 2016;Loveland et al., 2016), we do not detect a stable association of ΔRRM Rel with the ribosome upon a mupirocin challenge. This suggests that the interaction with the ribosome is significantly destabilised in Rel^ΔRRM^ and the protein dissociates during centrifugation. It is noteworthy that, despite an unstable association of Rel^ΔRRM^ with starved ribosomes, the expression of this protein strongly induces the accumulation of 100S ribosomal dimers, which is indicative of (p)ppGpp overproduction (Tagami et al., 2012). Note that the 100S formation is abrogated when the culture is treated with mupirocin (**Figure 4A** and **Supplementary Figure S5B**), most likely due to complete inhibition of translation by the antibiotic hindering expression of the 100S-promoting Hibernation Promoting Factor (HPF) which, in turn, is induced by accumulation of (p)ppGpp (Tagami et al., 2012).

**Figure 4.**
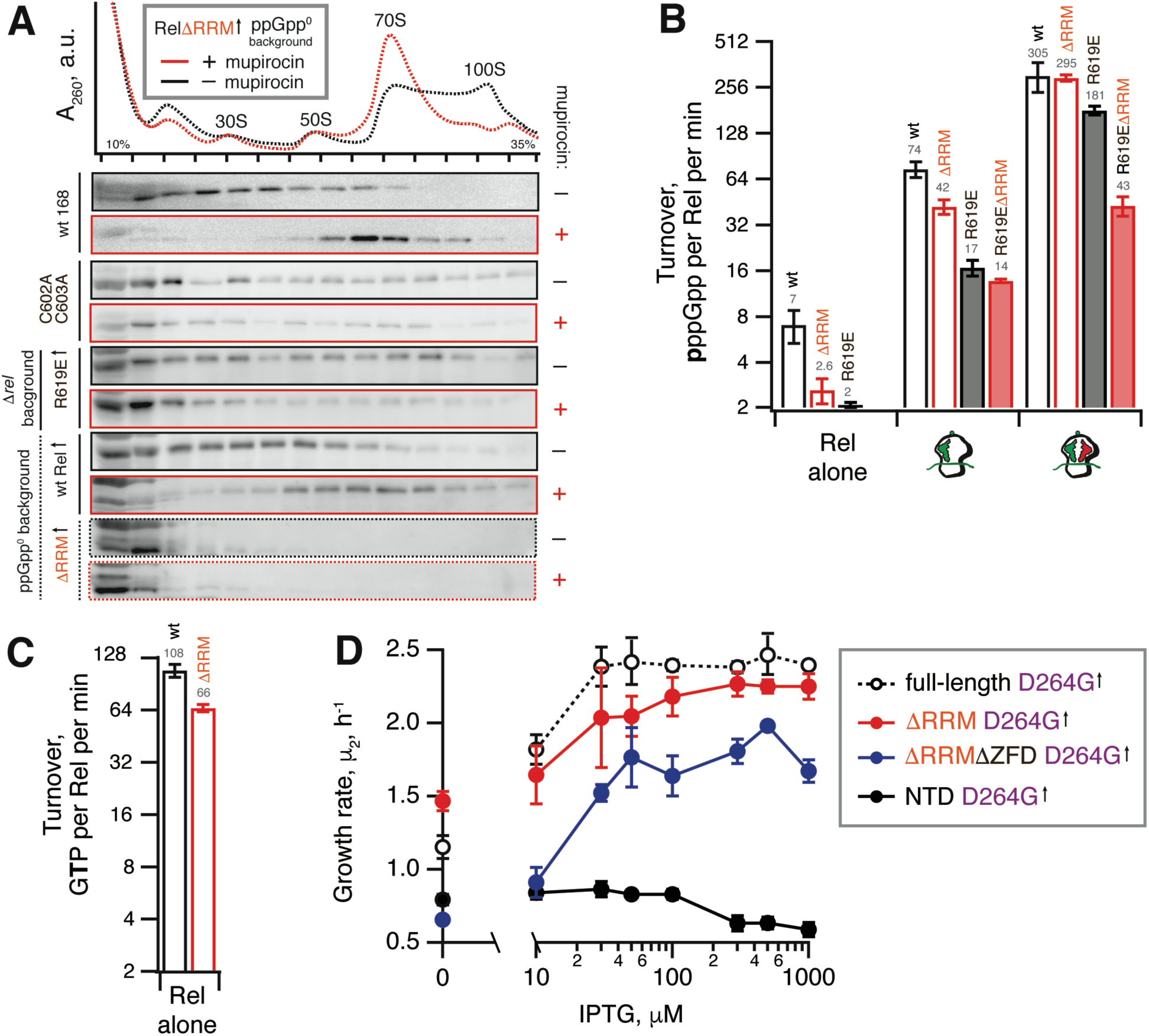
Deletion of the regulatory RRM domain destabilises Rel binding to starved ribosomal complexes and does not abrogate the hydrolysis activity. (**A**) Polysome profile and immunoblot analyses of Rel variants expressed either ectopically (↑) under the control of IPTG-inducible P*_hy-spnak_* promotor [ΔRRM Rel in ppGpp^0^ *B. subtilis* (VHB184), R619E Rel in Δ*rel B. subtilis* (VHB282)] or from the native chromosomal locus [C602A C603A mutant (VHB144)]. Expression of Rel^ΔRRM^ was induced by 1 mM IPTG for 10 minutes followed by a 10 minute challenge with 700 nM mupirocin. To drive the expression of R619E Rel, the strain as grown in LB supplemented with 1 mM IPTG. In the case of R619E and C602A C603A Rel the culture was treated with mupirocin for 20 minutes. Polysome profiles of all tested Rel variants are presented in **Supplementary Figure S5**, and an uncut version of a representative anti-Rel immunoblot is shown in **Supplementary Figure S2G**. (**B** and **C**) The effect of the RRM deletion on synthetic (**B**) and hydrolytic activity (**C**). The effects of the RRM deletion and the R619E mutation on Rel synthetic activity were assayed either alone or in the presence of either initiation and starved ribosomal complexes. The error bars represent standard deviations of the turnover estimates by linear regression using four data points. (**D**) The effects of titratable expression of synthesis-inactive D264G mutants (full-length VHB156, ΔRRM VHB162, ΔRRMΔZFD VHB163 and Rel^NTD^ VHB164) on Δ*rel B. subtilis* growing on liquid LB medium at 37 °C. The growth rates (µ^2^) were calculated from three independent biological replicates and the error bars represent standard deviations.

Our microbiological experiments demonstrate that both the C602A and C603A double substitution and the R619E point substitution render Rel^ΔRRM^ non-toxic (**Figure 3D**), which we attribute to further destabilisation of Rel’s interaction with starved ribosomal complexes. To directly probe the effects of these mutations, we used centrifugation experiments with full-length Rel carrying the substitutions. As expected, both the C602A C603A double mutant and R619E full-length variants are compromised in recruitment to the ribosome upon a mupirocin challenge (**Figure 4A**).

Taken together, these results demonstrate that Rel^ΔRRM^ is significantly more toxic than the full-length protein and this toxicity is dependent on the interaction with starved ribosomes, which is, in turn, destabilised in this truncation. As a next step, we set out to test the effects of RRM deletion – either alone or in combination with mutations further compromising the interactions with starved ribosomes – on Rel’s enzymatic activity in a reconstituted *B. subtilis* biochemical system.

### 3.5 Purification of RNA-free untagged *B. subtilis* Rel requires size-exclusion chromatography

To purify untagged *B. subtilis* Rel we combined our protocols used for purification of *E. coli* RelA (Turnbull et al., 2019) and *T. thermophilus* Rel (Van Nerom et al., 2019). (**Figure 5**). Importantly, during all of the chromatography steps we followed both absorbance at 260 and 280 nm complemented with SDS PAGE analysis of fractions. This is essential in order to identify and specifically pool the fractions containing Rel free from RNA contamination. After the initial capture using immobilized metal affinity chromatography (IMAC) in high ionic strength conditions (750 mM KCl) using HisTrap HP column charged with Zn^2+^ in order to avoid possible replacement of the Zn^2+^ in the ZFD domain by Ni^2+^ ions (Block et al., 2009), His_10_-SUMO-Rel was applied on size-exclusion chromatography (SEC) on HiLoad 16/600 Superdex 200 pg column (**Figure 5D**). Both in the case of the full length (**Figure 5B**) and ΔRRM Rel (**Figure 5C**), the RNA-free fractions constitute the minority of the protein that elute considerably later than the bulk of the RNA-contaminated Rel. While the SEC step is essential for generating RNA-free Rel preparations, the majority of the protein prep is lost at this stage. After the SEC step, the buffer was exchanged to storage buffer containing arginine and glutamic acid that improve protein solubility and long-term stability (Golovanov et al., 2004) (**Figure 5D**), His_10_-SUMO tag was cleaved off and removed by passing the protein via second IMAC (**Figure 5E**). The quality of the final preparations was assessed by SDS-PAGE (**Figure 5F** and **Supplementary Figure S6**) as well as spectrophotometrically: OD_260_/OD_280_ ratio below 0.8 corresponding to less than 5% RNA contamination (Layne, 1957).

**Figure 5.**
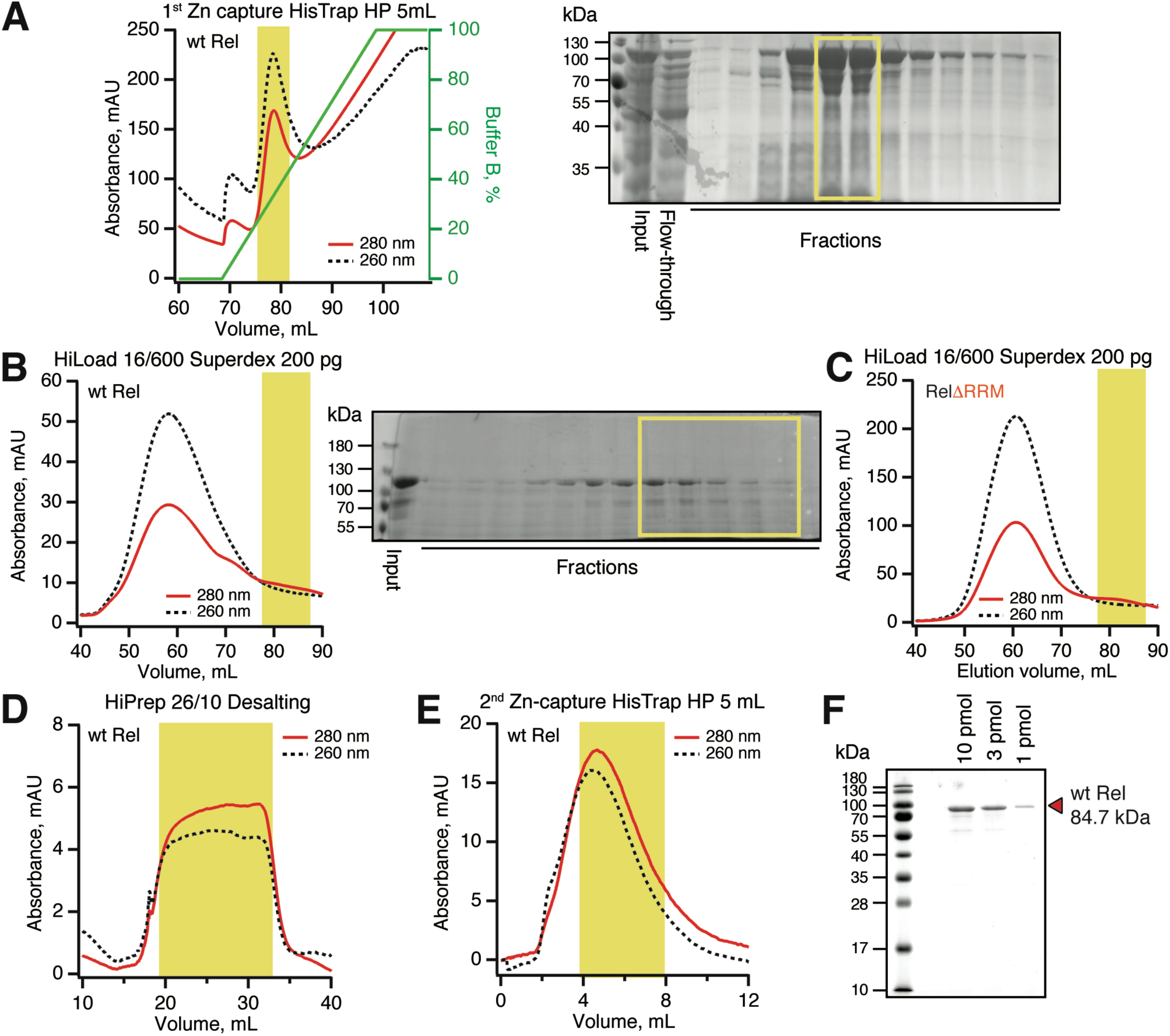
Purification of RNA-free untagged *B. subtilis* Rel. N-terminally His_10_-SUMO tagged RelA was overexpressed and purified as described in detail in *Materials and Methods*. (**A**) Cells were lysed and subjected to immobilized metal affinity chromatography (IMAC) using a Zn^2+^-charged HisTrap 5 mL HP column. The fraction corresponding to Rel with the lowest contamination of nucleic acids (highlighted in yellow) was carried forward. Size-exclusion chromatography on HiLoad 16/600 Superdex 200 pg was used to further separate the RNA-free Rel fractions (**B**: full-length wild type Rel; **C**: ΔRRM Rel). Following the buffer exchange on HiPrep 10/26 desalting column (**D**), the boxed-out fractions were pooled and the His_10_-SUMO tag was cleaved off by the His_6_-Ulp1 protease. (**E**) Native untagged Rel was separated from His_6_-Ulp1 and the His_10_-SUMO tag by the second round of IMAC. Highlighted fractions were pooled, concentrated, aliquoted, flash-frozen in liquid nitrogen and stored at –80 °C. (**F**) SDS-PAGE analysis of the purified native untagged *B. subtilis* Rel.

We have tested the effects of omission of the SEC step on the purification and activity of *B. subtilis* Rel preparations. Without the SEC step, the OD_260_/OD_280_ ratio was dramatically higher (1.9), suggesting that, counterintuitively, the ‘no SEC’ Rel preparation predominantly contains not protein but RNA. We have resolved the sample on 15% SDS-PAGE (**Figure 6A**) and denaturating 1.2% agarose (2% formaldehyde) (**Figure 6B**) gels, as well as subjected the samples to negative staining electron microscopy (**Figure 6C**). While the SDS-PAGE gel revealed multiple protein bands with Mw between 40 and 10 kDa, the agarose gel revealed that the RNA contaminant is dominated by three distinct populations of approximately 3000, 1500 and 100 nucleotides in length. Large (approximately 20 nm in diameter) particles are clearly visible on the to negative staining EM images. Collectively, this suggests that the RNA contamination is dominated by ribosomal particles, although it is unclear whether these are intact or partially degraded. Taking into account that 1 A_260_ corresponds to 23 pmol 70S particles, we estimate that our ‘no SEC’ preparations contain 45 nM 70S ribosomes per 1 µM Rel, which corresponds to sub-stoichiometric contamination of 5 % of Rel being in complex, and 95% free. We next tested the effects of SEC omission on the enzymatic activity of Rel. The effects are exceedingly mild. The synthetase activity is virtually unaffected; importantly, activation by deacylated tRNA remains strictly mRNA-dependent, with tRNA^Val^ inducing the enzymatic activity of ‘no SEC’ Rel only in the presence of 70S initiation complexes (70S IC) but not vacant 70S ribosomes (**Figure 6D**). Importantly, since ribosomes or starved complexes are added in our synthetase assays in excess over Rel (500 nM vs 140 nM Rel), in the final reaction mixture purified ribosomes are in approximately 100x excess over the contaminant. Finally, the hydrolase activity of ‘no SEC’ Rel is approximately two-fold lower (**Figure 6E**), which, however, could reflect variability in preparations.

**Figure 6.**
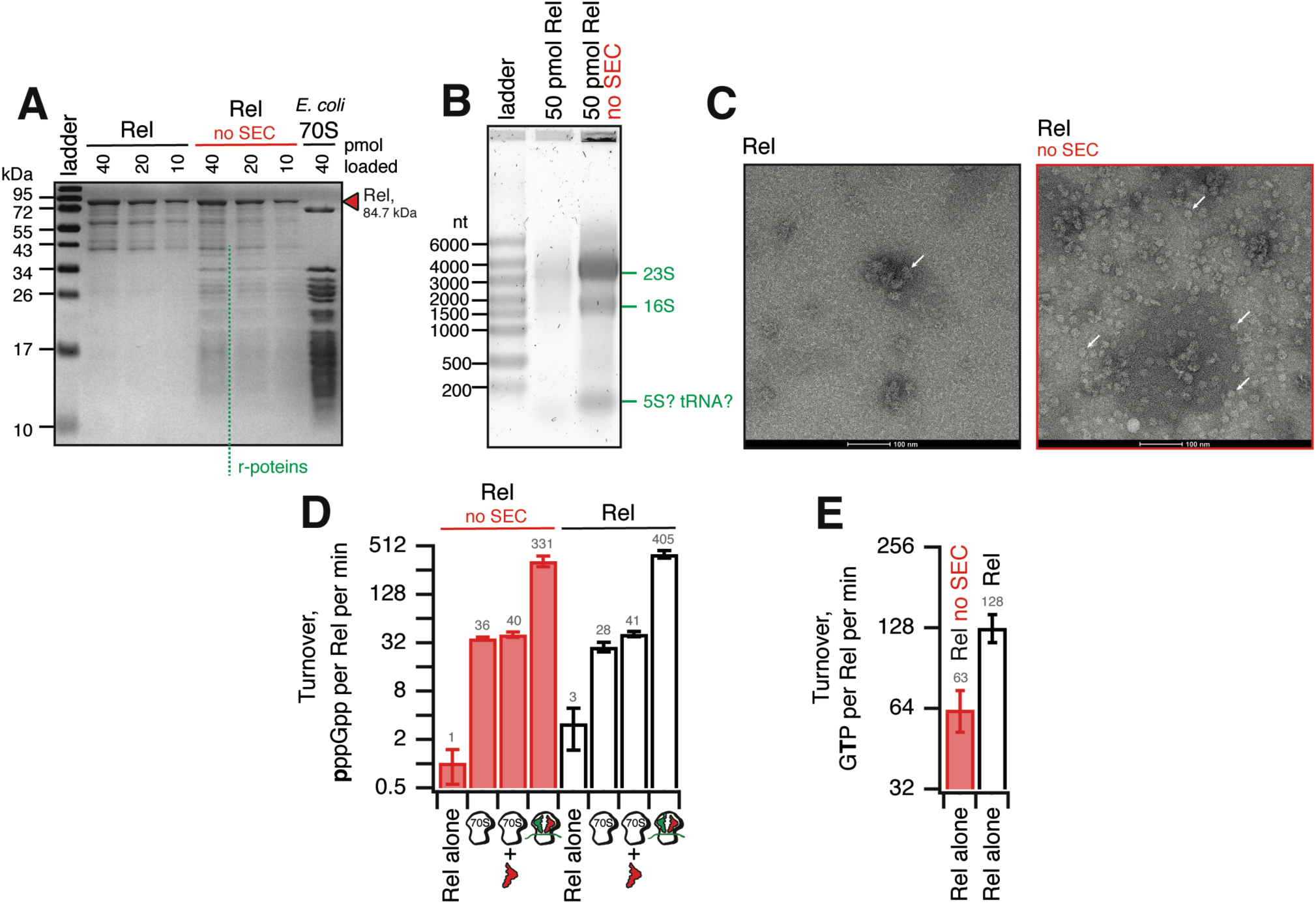
Omission of the size-exclusion chromatography step results in sub-stoichiometric contamination of *B. subtilis* Rel preparations with *E. coli* ribosomal particles. (**A**) SDS-PAGE analysis of wild-type full-length Rel protein purified either as described in Figure 5 or with the SEC step omitted (no SEC). Desaturating agarose gel (**B**) and negative staining electron microscopy (**C**) analyses of full-length Rel protein purified with and without the SEC step. Individual ribosomal particles are indicated with white arrows on (**C**). Effects of the SEC omission on synthetase (**D**) and hydrolase (**E**) activity of *B. subtilis* Rel. The synthetase activity was assayed with either Rel alone or in the presence of 0.5 µM 70S, 70S supplemented with 2 µM deacylated tRNA^Val^ and no mRNA or starved ribosomal complexes (0.5 µM 70S IC(MVF) supplemented with 2 µM deacylated tRNA^Val^). The error bars represent standard deviations of the turnover estimates by linear regression using four data points.

Taken together, these results suggest that in the absence of a dedicated SEC step, Rel preparations are sub-stoichiometrically contaminated with ribosomes. While the effects of this contamination on the enzymatic activities of Rel are minor, it might interfere with other assays (see *Discussion*).

### 3.6 The R619E ZFD substitution compromises activation of *B. subtilis* Rel^ΔRRM^ by starved ribosomal complexes

We tested the ^3^H-pppGpp synthesis by full-length Rel as well as Rel^ΔRRM^, either alone or activated by the ribosomes or starved complexes in a reconstituted system (**Figure 4B**). When the protein is tested by itself, the Rel^ΔRRM^ mutant is less active than the full-length, suggesting that deletion of the RRM domain does not lead to the loss of auto-inhibition. While Rel^ΔRRM^ remains less active than the full-length when activated by initiation complexes (about two-fold), in the presence of starved ribosomal complexes the two proteins are equally active. This could be explained by tRNA stabilising Rel on the ribosome and overriding the defect caused by the removal of the RRM domain.

The R619E mutation compromises activation of the full-length Rel by the initiation complexes (more than four-fold), and the effect is less pronounced in the presence of deacylated tRNA^Val^ (less than two-fold decrease in activity). Just as in the case of the RRM deletion, a possible explanation is that the deacylated tRNA strongly stimulates the binding of Rel to the ribosome and offsets the effect of the mutation R619E. When the R619E substitution is introduced into ΔRRM Rel, the combination of the two mutations destabilising Rel binding to the ribosome results in compromised activation both by the initiation (five-fold) and starved (seven-fold) ribosomal complex. Despite several attempts we failed to generate sufficiently pure and soluble C602A C603A Rel^ΔRRM^, which precluded direct biochemical characterisation of this mutant.

Taken together, our biochemical results demonstrate that while RRM is important in Rel recruitment to the ribosome, this domain is not absolutely essential for the activation of its (p)ppGpp synthesis activity by starved ribosomal complexes, which is consistent with the ribosome-dependent nature of the Rel^ΔRRM^ toxicity in live cells.

### 3.7 The RRM deletion moderately decreases the hydrolysis activity of *B. subtilis* Rel

It was recently proposed that the RRM domain has a stimulatory effect on the hydrolysis activity of *B*. *crescentus* Rel, and the loss of this regulatory mechanism explains the toxicity of the ΔRRM mutant (Ronneau et al., 2019). This hypothesis does not explain the toxicity of the ΔRRM variant of the synthesis-only RSH RelA (**Figure 2** and (Turnbull et al., 2019)) and our microbiological experiments showing that the synthesis-defective ΔRRM D264G variant of *B. subtilis* Rel remains active as a (p)ppGpp hydrolase (**Figure 3A**). Importantly, the variant characterised by Ronneau and colleagues (*C. crescentus* Rel^Δ668-719^) does not completely lack the RRM, and it is possible that the remaining beta-strand alpha-helix turn structural element was interfering with the hydrolysis activity of the construct (Ronneau et al., 2019).

Our enzymatic assays following ^3^H-pppGpp degradation by full-length and Rel^ΔRRM^ show that the latter is approximately twice less active (**Figure 4C**). While the defect is detectable, it is quite minor. To test if the hydrolysis defect is more pronounced in the living cell, we re-tested the hydrolysis activity of synthesis-deficient SYNTH D264G full-length Rel as well as C-terminally truncated Rel variants in the Δ*rel* background using the growth rate (µ_2_) as a proxy (**Figure 4D**). We modulated the expression levels by titrating the inducer, IPTG, from 10 to 1000 µM. In good agreement with the biochemical results demonstrating a minor defect in hydrolysis caused by deletion of the RRM domain, ΔRRM D264G Rel mutant promotes the growth of Δ*rel B. subtilis* only moderately less efficiently than the full-length D264G.

### 3.8 The hydrolysis activity of *B. subtilis* Rel is not activated by branched-chain amino acids binding to the RRM domain

It was also recently reported that binding of branched-chain amino acids (BCAAs) to the ACT/RRM domain induces the hydrolysis activity of *Rhodobacter capsulatus* Rel (Fang and Bauer, 2018). The N651A substitution abrogates amino acid binding to *R. capsulatus* Rel^CTD^ and leads to (p)ppGpp accumulation in the cell, presumably due to lower hydrolysis activity of the mutant enzyme. It is, therefore, conceivable that *B. subtilis* Rel^ΔRRM^ is less hydrolytically active in the cell than the full-length protein due to the loss of BCAA-mediated activation. While the *B. subtilis* Rel^CTD^ fragment was shown to preferentially bind leucine with a *K_D_* of 225 µM, no enzymatic assays were performed with this protein (Fang and Bauer, 2018). Notably, while the CTD region of *E. coli* RelA binds valine with high affinity (*K_D_* of 2.85 µM) (Fang and Bauer, 2018), this interaction could not be regulating the hydrolysis activity of this synthesis-only RSH. It is, therefore, unclear whether *B. subtilis* Rel is, indeed, regulated by branched-chain amino acids similarly to *R. capsulatus* enzyme. Therefore, we tested the effect of 1 mM leucine on ^3^H-pppGpp degradation by *B. subtilis* Rel. We detect no stimulatory effect (**Figure 7A**). Furthermore, when we introduced the N685A substitution (equivalent to N651A in *R. capsulatus*) in the chromosomal *rel* gene, we detected no growth defect in comparison to wild-type 168 *B. subtilis*, either on solid or liquid LB media (**Figure 7B-D**). Taken together, these results suggest that amino acid binding to RRM should not automatically be equated with regulation of the hydrolysis activity.

**Figure 7.**
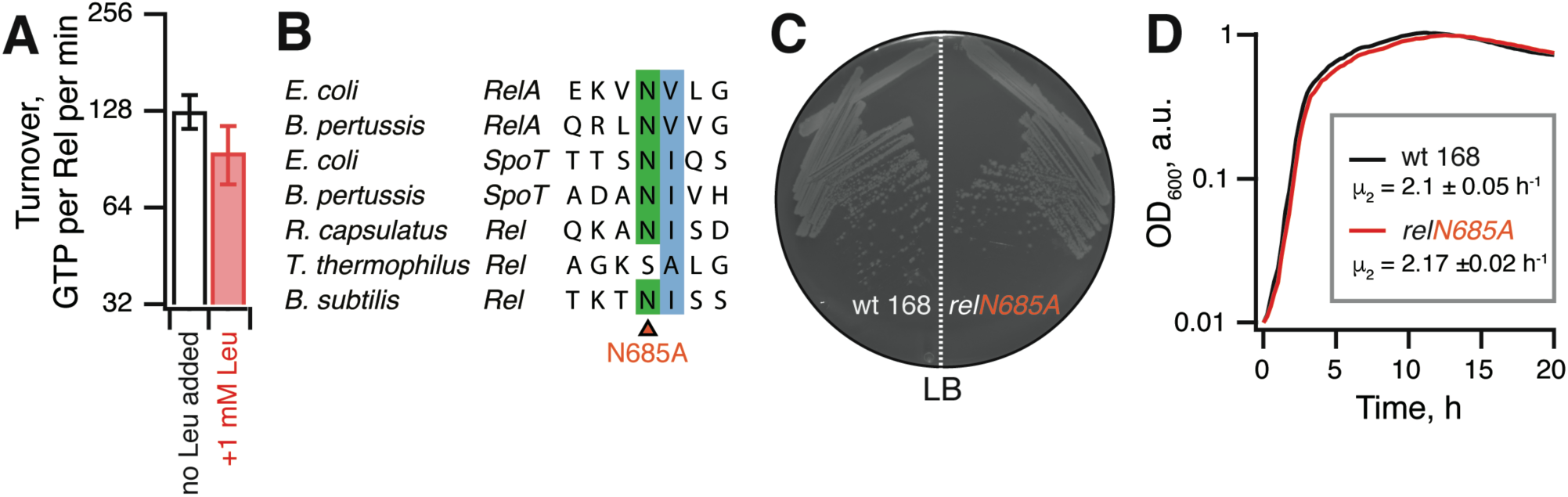
The hydrolysis activity of *B. subtilis* Rel is not activated by branched-chain amino acids binding to the RRM domain. (**A**) Hydrolase activity of *B. subtilis* Rel is not stimulated by leucine. The experiments were performed in HEPES:Polymix buffer, pH 7.5 at 37 °C in the presence of 5 mM Mg^2+^. Error bars represent standard deviations of the turnover estimates by linear regression using four data points. (**B**) Sequence alignment of the *B. subtilis* N685 region of representative long RSHs. (**C** and **D**) Wild-type *B. subtilis* 168 and isogenic chromosomal *relN685A* mutant (VHB455) were grown on solid (**C**) and liquid (**D**) LB media at 37 °C. The plate was scored after 12 hours at 37 °C. The growth rates (µ^2^) were calculated from three independent biological replicates and the error bars (shown as shadows) represent standard deviations. Since the liquid media growth experiments were performed in plates format in Bioscreen C growth curve analysis system the OD_600_ is presented in arbitrary units.

## 4 Discussion

Taken together, our results demonstrate that i) deletion of the RRM domain renders *B. subtilis* Rel and *coli* RelA toxic due to (p)ppGpp overproduction in a ribosome and tRNA-dependent manner, ii) RRM deletion does not abrogate the (p)ppGpp hydrolysis activity of *B. subtilis* Rel, iii) RRM deletion destabilises the interaction of *B. subtilis* Rel with starved ribosomal complexes, and iv) this destabilisation renders the mutant enzyme more sensitive than the full length Rel to deactivation by additional substitutions further compromising its association with starved ribosomes. Our biochemical results do not explain why exactly ΔRRM Rel/RelA is toxic: *B. subtilis* Rel^ΔRRM^ behaves as a weaker binder of starved ribosomal complexes (**Figure 4A**) and is less enzymatically active in (p)ppGpp synthesis assays (**Figure 4BC**) as compared to the full-length Rel. A dedicated follow-up study is necessary to clarify this question. *E. coli* RelA^ΔRRM^ displays a similar – although more pronounced – defect in activation by starved ribosomal complexes (Takada et al., 2020).

Our report expands the mutation toolbox for dissecting the molecular mechanisms of long RSH enzymes. We confirm that, as was shown for H432E *E. coli* RelA mutant (Winther et al., 2018), the corresponding H420E mutation in *B. subtilis* Rel is a useful tool for specifically abrogating activation of Rel by starved ribosomal complexes. Additionally, we demonstrate the utility of two novel mutations in the ZFD domain: R619E (R629E in *E. coli* RelA) as well as a double C602A C603A substitution (C612A C613A in *E. coli* RelA). While these mutations do not completely abrogate the activation, acting in epistasis they can be employed to reveal weaker phenotypes or effects, as was shown in the current work when the mutations were combined with RRM/ACT deletion.

Finally, we would like to draw the attention of the research community working on long RSH enzymes to technical aspects of protein purification. It is common to purify Rel/RelA for biochemical experiments using a single-step purification (for example (Gratani et al., 2018;Wood et al., 2019)). However, both RelA (Turnbull et al., 2019) and Rel have a strong tendency for RNA contamination and multiple additional steps are necessary to remove this contaminant. Therefore, reporting the 260/280 absorbance ratio of the final preparations is essential. RNA-free protein preparations typically have a 260/280 absorbance ratio of 0.57, but this parameter can vary depending on the amino acid composition, specifically the abundance of tryptophan and phenylalanine (Layne, 1957). Without extra steps to remove RNA contamination, single-step preparations are likely to be heavily contaminated with ribosomal particles (**Figure 6**), which is likely to interfere with the estimation of the oligomerisation state of Rel, since the RNA-bound protein elutes much earlier than the RNA-free fraction (**Figure 5BC**). This contamination may explain the surprising observation that the addition of Ni-NTA purified *S. aureus* Rel inhibits the 50S assembly factor DEAD-box RNA helicase CshA (Wood et al., 2019). The unlabelled contaminating ribosomal particles could potentially be recognised by CshA, thus acting as a competitor in the helicase assay that uses a synthetic Cy3-labelled RNA duplex as a substrate. It is also plausible that formation of stable complexes of Rel with ribosomes in live *E. coli* could generate false-positive signals in bacterial two-hybrid assays, accounting for the observed protein-protein interaction between *S. aureus* Rel and the ribosome assembly factors Era and CshA (Wood et al., 2019).

## 7 Author Contributions

HT conceived the study, HT, MR and VH designed experiments, HT performed experiments with *B. subtilis* Rel, ID and MR performed experiments with *E. coli* RelA, VM performed electron microscopy, RM and GA contributed tools and reagents, GCA and AG-P performed structure and sequence analyses, HT and VH drafted and revised the manuscript with contributions from MR, AG-P and GCA. All authors have read and approved the final version of this manuscript.

## Acknowledgments

We would like to thank the Protein Expertise Platform (PEP) at Umeå University and Mikael Lindberg for constructing pET24d::*His_10_-SUMO*-based expression constructs and purifying His_6_-Ulp1, and Julien Caballero-Montes for comments on the manuscript. The electron microscopy data was collected at the Umeå Core Facility for Electron Microscopy, a node of the Cryo-EM Swedish National Facility, funded by the Knut and Alice Wallenberg, Family Erling Persson and Kempe Foundations, SciLifeLab, Stockholm University and Umeå University.

