## Supplementary Material for "The C-terminal RRM/ACT domain is crucial for fine-tuning the activation of ‘long’ RelA-SpoT Homolog enzymes by ribosomal complexes"

### 1.1 Construction of *E. coli* plasmids

Mutations in RelA$^{\Delta RRM}$ ORF were introduced using two step PCR (Ho et al., 1989). Specifically, primers MR3 and MR4 were used to introduce H432E, MR5 and MR6 to introduce R629E, MR7 and MR8 to introduce C612A/C613A; primers MR1 (forward) and MR2 (reverse) were universally used to amplify the PCR products. The resultant linear fragments were digested with *Bam*HI and *Xho*I enzymes (ThermoScientific) and cloned into pNDM220 (Gotfredsen and Gerdes, 1998) plasmid cut with the same restriction enzymes.

### 1.2 Construction of *B. subtilis* strains and plasmids

To construct the strain NBS1437 [*trpC2 relQ::tet relP::spcR*], first, two ≈500 nucleotide-long DNA fragments were amplified by PCR using wt168 genomic DNA as a template: one located upstream of the *relQ* ORF (primers: NOP7 and NOP8), another located downstream of the *relQ* ORF (primer, NOP11 and NOP12). Second, the tetracycline resistance marker (*tet*) of pDG1515 (Guerout-Fleury et al., 1995) was amplified by PCR using the primers NOP9 and NOP10. All three PCR products described above were used simultaneously as the template for PCR amplification using primers NOP7 and NOP12. The resulting long fragment was used to transform strain 168 (*trpC2*). *relQ* deletion mutants were selected by tetracycline resistance, yielding the NBS1393 [*trpC2 relQ::tet*] strain. Chromosomal DNA extracted from strain RIK908 [*trpC2 relP::spcR*] was used to transform the strain NBS1393 and selected by spectinomycin resistance yielding the strain NBS1437.

To construct the strain VHB63 [*trpC2 ΔrelQ relP::cmR rel:::spcR*] strain, first, two ≈2500 nucleotide-long DNA fragments were amplified by PCR using wt168 genomic DNA as a template: one located upstream of the *rel* ORF (primers: VHT29 and NOP13), another located downstream of the *rel* ORF (primers: NOP16 and VHT30). Second, spectinomycin resistant gene (*spcR*) of pDG1727 (Guerout-Fleury et al., 1995) was amplified by PCR using the primers NOP14 and NOP15. All three PCR products described above were used simultaneously as the template for PCR amplification using primers VHT29 and VHT30. The resulting long fragment was used to transform strain 168 [*trpC2*]. *rel* deletion mutants were selected for spectinomycin resistance, yielding the NHT437 [*trpC2 rel::spcR*] strain. Chromosomal DNA extracted from strain NHT437 was used to transform the strain NHT436 [*trpC2 ΔrelQ relP::cmR*] and selected for spectinomycin resistance yielding the VHB63 strain.

To construct the VHB47 strain [*trpC2 rplK::cmR*], first, two ≈1500 nucleotide-long DNA fragments were amplified by PCR using wt168 genomic DNA as a template: one located upstream of the *rplK* ORF (primers: VHT1 and VHT5), another located downstream of the *rplK* ORF (primers: VHT2 and VHT9). Second, the chloramphenicol-resistance marker (*cmR*) of pC194 (Horinouchi and Weisblum, 1982) was amplified by PCR using the primers VHT6 and VHT10. All three PCR products described above were used simultaneously as the template for PCR amplification using primers VHT1 and VHT2. The resulting long fragment was used to transform strain 168 (*trpC2*). *rplK* deletion mutants were selected by chloramphenicol resistance, yielding the strain VHB47. Chromosomal DNA extracted from strain VHB47 was used to transform the strain NBS1437 [*trpC2 relQ::tet relP::spcR*], and selected for chloramphenicol resistance, yielding the strain VHB49 [*trpC2 relQ::tet relP::spcR rplK::cmR*].

To construct the strain NBS1729 [*trpC2 rel-spcR*], First, two ≈1000 nt-long DNA fragments were amplified by PCR using genomic DNA as a template: one located +1452 ~ +2205 relative to the translation initiation codon of the rel *rel* ORF (primers: NOP1 and NOP2), another located downstream of the *rel* ORF (primers: NOP5 and NOP6). Second, the spectinomycin resistant gene (*spcR*) of pDG1727 (Guerout-Fleury et al., 1995) was amplified by PCR using the primers NOP3 and NOP4. All three PCR products described above were used simultaneously as the template for PCR amplification using primers NOP1 and NOP6. The resulting long fragment was used to transform strain 168 (*trpC2*). Transformants were selected for spectinomycin resistance yielding the strain NBS1729.

Strain VHB68 [*trpC2 relH420E-spcR*] was constructed using the site-directed mutagenesis technique (Heckman and Pease, 2007). To exchange the CAC codon (coding for Histidine) at position 420 for AGA (coding for Glutamic acid), two ≈2500 nucleotide-long overlapping DNA fragments were amplified by PCR using genomic DNA of strain NBS1729 as a template. The first fragment used the region located upstream of the *rel* ORF as the DNA template, spanning to codon 420 (primer, VHT29 and VHT32). The second used the region starting with the codon 420 as a template, spanning downstream of the *rel* ORF to include the *spcR* resistance marker (primers: VHT30 and VHT31). Two PCR products described above were used simultaneously as the template for PCR amplification using primers VHT29 and VHT30. The resulting long fragment was used to transform strain 168 (*trpC2*). The transformant was selected for spectinomycin resistance, yielding the strain VHB68. The desired mutation was confirmed by sequencing. Chromosomal DNA extracted from strain VHB68 was used to transform the strain NHT436, and bacteria were selected for spectinomycin resistance, yielding the strain VHB60 [*trpC2 ΔrelQ relP::cmR relH420E-spcR*].

Strain VHB144 [*trpC2 relC602C603-spcR*] was constructed using the site-directed mutagenesis technique. To exchange the TGC codon (coding for Cysteine) at position 602 and 603 for GCC (coding for Alanine), two ≈2500 nucleotide-long overlapping DNA fragments were amplified by PCR using the genomic DNA of strain NBS1729 strain as a template. The first fragment used the region located upstream of the *rel* ORF as the DNA template, spanning to codon 603 (primers: VHT29 and VHT40). The second used as a template the region starting with the codon 602 of *rel* ORF, spanning downstream of the *rel* ORF to include the *spcR* resistance marker (primers: VHT30 and VHT39). The two PCR products described above were used simultaneously as the template for PCR amplification using primers VHT29 and VHT30. The resulting long fragment was used to transform strain 168 (*trpC2*). The transformant was selected by spectinomycin resistance yielding the strain VHB144. The desired mutation was confirmed by sequencing. Chromosomal DNA extracted from strain VHB144 was used to transform the strain NHT436, and selected for spectinomycin resistance, yielding the strain VHB62 [*trpC2 ΔrelQ relP::cmR relC602AC603A-spcR*].

To construct the strain VHB155 [*trpC2 amyE::*$P_{hy\text{-}spank}$*-rel cmR rel::ermR*], the region containing *rel* ORF as well as the upstream Shine-Dalgarno sequence was amplified by PCR using *B. subtilis* 168 wild-type genomic DNA as a template (primers: NSP1 and NSP2). The PCR product was digested with SalI and SphI restriction enzymes and inserted into the SalI and SphI sites of plasmids pHT001 encoding i) a chloramphenicol-resistance marker (*cmR*), ii) a polylinker downstream of isopropyl-β-D-thiogalactopyranoside (IPTG) inducible $P_{hy\text{-}spank}$ promoter and iii) the lac repressor ORF (all three inserted in the middle of the *amyE* gene) (Takada et al., 2014), yielding the plasmid pHT101 (pHT001-*rel*, containing *rel* gene whose transcription is controlled by $P_{hy\text{-}spank}$ promoter) which was used to transform wild type 168 strain. Selection for chloramphenicol resistance yielded the VHB145 [*trpC2 amyE::*$P_{hy\text{-}spank}$*-rel cmR*]. DNA extracted from strain RIK900 (*trpC2 rel::ermR*) was used to transform the strain VHB145, and selected by erythromycin resistance yielding the desired strain VHB155. To construct the strain VHB183 [*trpC2 relQ::tet relP::spcR amyE*::$P_{hy\text{-}spnak}$*-rel cmR rel::ermR*], plasmid pHT101 was used to transform strain NBS1437, and selected for chloramphenicol resistance, yielding the strain VHB175 [*trpC2 relQ::tet relP::spcR amyE*::$P_{hy\text{-}spnak}$*-rel cmR*]. DNA extracted from strain RIK900 was used to transform the strain VHB175, and selected for erythromycin resistance, yielding the desired strain VHB183.

To construct the strain VHB156 [*trpC2 amyE::*$P_{hy\text{-}spank}$*-relD264G cmR rel::ermR*], plasmid pHT101 was subjected to site-directed mutagenesis using primer NHP1 and NHP2 , according to directions of Phusion Site-Directed Mutagenesis Kit (Thermo Fisher Scientific), yielding the plasmid pHT125 (pHT001-*relD264G*) which was used to transform wt168 strain. Selection for chloramphenicol resistance yielded the strain VHB151 [*trpC2 amyE::*$P_{hy\text{-}spank}$*-relD264G cmR*]. DNA extracted from strain RIK900 was used to transform the strain VHB145, and selected for erythromycin resistance, yielding the desired strain VHB156. To construct the strain VHB187 [*trpC2 relQ::tet relP::spcR amyE*::$P_{hy\text{-}spnak}$*-relD264G cmR rel::ermR*], plasmid pHT125was used to transform the strain NBS1437, and selected for chloramphenicol resistance, yielding the strain VHB179 [*trpC2 relQ::tet relP::spcR amyE*::$P_{hy\text{-}spnak}$*-relD264G cmR*]. DNA extracted from strain RIK900 was used to transform the strain VHB179, and selected by erythromycin resistance, yielding the desired strain VHB187.

To construct the strain VHB159 [*trpC2 amyE::*$P_{hy\text{-}spank}$*-rel* $^{\Delta RRM}$ *cmR rel::ermR*], plasmid pHT101 was subjected to site-directed mutagenesis using primer VHT68 and VHT69 , according to directions of Phusion Site-Directed Mutagenesis Kit (Thermo Fisher Scientific), yielding the plasmid VHP435 (pHT001- *rel* $^{\Delta RRM}$), which was used to transform wild type 168 strain. Selection for chloramphenicol resistance yielded the strain VHB146 [*trpC2 amyE::*$P_{hy\text{-}spank}$*-rel* $^{\Delta RRM}$ *cmR*]. DNA extracted from strain RIK900was used to transform the strain VHB146, and selected by erythromycin resistance, yielding the desired strain VHB159. To construct the strain VHB184 [*trpC2 relQ::tet relP::spcR amyE*::$P_{hy\text{-}spnak}$*-rel* $^{\Delta RRM}$ *cmR rel::ermR*], plasmid VHP435 was used to transform the strain NBS1437, and selected by chloramphenicol resistance, yielding the strain VHB176 [*trpC2 relQ::tet relP::spcR amyE*::$P_{hy\text{-}spnak}$*-rel* $^{\Delta RRM}$ *cmR*]. The genomic DNA of the RIK900 strain was used to transform the strain VHB176, after selection for erythromycin resistance yielding the desired strain VHB184.

To construct the strain VHB160 [*trpC2 amyE::*$P_{hy\text{-}spank}$*-rel* $^{\Delta ZFD\text{-}RRM}$ *cmR rel::ermR*], plasmid pHT101 was subjected to site-directed mutagenesis using primer VHT69 and VHT189, according to directions of Phusion Site-Directed Mutagenesis Kit (Thermo Fisher Scientific), yielding VHP434 (pHT001- *rel* $^{\Delta ZFD\text{-}RRM}$) plasmid which was used to transform wt168 strain. Selection for chloramphenicol resistance yielded the strain VHB147 [*trpC2 amyE::*$P_{hy\text{-}spank}$*-rel* $^{\Delta ZFD\text{-}RRM}$ *cmR*]. The genomic DNA of the RIK900 strain was used to transform the strain VHB147, after selection for erythromycin resistance yielding the desired strain VHB160. To construct the strain VHB185 [*trpC2 relQ::tet relP::spcR amyE*::$P_{hy\text{-}spnak}$*-rel* $^{\Delta ZFD\text{-}RRM}$ *cmR rel::ermR*], plasmid VHP434 was used to transform strain NBS1437, and selected by chloramphenicol resistance, yielding the strain VHB177 [*trpC2 relQ::tet relP::spcR amyE*::$P_{hy\text{-}spnak}$*-rel* $^{\Delta ZFD\text{-}RRM}$ *cmR*]. The genomic DNA of the RIK900 strain was used to transform the strain VHB177, and selected by erythromycin resistance, yielding the desired strain VHB185.

To construct the strain VHB161 [*trpC2 amyE::*$P_{hy-spank}$*-rel(1-373)* $^{\Delta CTD}$ *cmR rel::ermR*], the pHT101 plasmid was subjected to site-directed mutagenesis using primers VHT69 and VHT190, as per manufacturers manual (Phusion Site-Directed Mutagenesis Kit, Thermo Fisher Scientific), yielding the plasmid VHP433 (pHT001-*rel(1-373)* $^{\Delta CTD}$) which was used to transform wt168 strain. Selection for chloramphenicol resistance yielded the strain VHB148 [*trpC2 amyE::*$P_{hy-spank}$*-rel(1-373)* $^{\Delta CTD}$ *cmR*]. DNA extracted from strain RIK900 was used to transform the strain VHB148, and selected by erythromycin resistance, yielding the desired strain VHB161. To construct the strain VHB186 [*trpC2 relQ::tet relP::spcR amyE*::$P_{hy-spnak}$-*rel(1-373)* $^{\Delta CTD}$ *cmR rel::ermR*], plasmid VHP433 was used to transform strain NBS1437, and selected by chloramphenicol resistance, yielding the strain VHB178 [*trpC2 relQ::tet relP::spcR amyE*::$P_{hy-spnak}$-*rel(1-373)* $^{\Delta CTD}$ *cmR*]. The genomic DNA of the RIK900 strain was used to transform the strain VHB178, and selected by erythromycin resistance, yielding the desired strain VHB186.

To construct the strain VHB162 [*trpC2 amyE::*$P_{hy-spank}$*-reD264G* $^{\Delta RRM}$ *cmR rel::ermR*], plasmid pHT125 was subjected to site-directed mutagenesis using primers VHT68 and VHT69, according to directions of Phusion Site-Directed Mutagenesis Kit (Thermo Fisher Scientific), yielding plasmid VHP438 (pHT001-*relD264G*$^{\Delta RRM}$) which was used to transform wt168 strain. Selection for chloramphenicol resistance yielded the strain VHB152 [*trpC2 amyE::*$P_{hy-spank}$*-relD264G* $^{\Delta RRM}$ *cmR*]. DNA extracted from strain RIK900 (*trpC2 rel::ermR*) was used to transform the strain VHB152, and selected by erythromycin resistance, yielding the desired strain VHB162. To construct the strain VHB188 [*trpC2 relQ::tet relP::spcR amyE*::$P_{hy-spnak}$-*relD264G* $^{\Delta RRM}$ *cmR rel::ermR R*], plasmid VHP438 was used to transform strain NBS1437, and selected by chloramphenicol resistance, yielding the strain VHB180 [*trpC2 relQ::tet relP::spcR amyE*::$P_{hy-spnak}$-*relD264G* $^{\Delta RRM}$ *cmR*]. The genomic DNA of the RIK900 strain was used to transform the strain VHB180, and selected by erythromycin resistance, yielding the desired strain VHB188.

To construct the strain VHB163 [*trpC2 amyE::*$P_{hy-spank}$*-relD264G* $^{\Delta ZFD-RRM}$ *cmR rel::ermR*], plasmid pHT125 was subjected to site-directed mutagenesis using primers VHT69 and VHT189, as per manufacturers manual (Phusion Site-Directed Mutagenesis Kit, Thermo Fisher Scientific), yielding the VHP437 plasmid (pHT001-*relD264G* $^{\Delta ZFD-RRM}$) which was used to transform wt168 strain. Selection for chloramphenicol resistance yielded the strain VHB153 [*trpC2 amyE::*$P_{hy-spank}$*-relD264G* $^{\Delta ZFD-RRM}$ *cmR*]. DNA extracted from strain RIK900 was used to transform the strain VHB153, and selected by erythromycin resistance, yielding the desired strain VHB163. To construct the strain VHB189 [*trpC2 relQ::tet relP::spcR amyE*::$P_{hy-spnak}$-*relD264G* $^{\Delta ZFD-RRM}$ *cmR rel::ermR R*], plasmid VHP437 was used to transform strain NBS1437, and selected by chloramphenicol resistance, yielding the strain VHB181 [*trpC2 relQ::tet relP::spcR amyE*::$P_{hy-spnak}$-*relD264G* $^{\Delta ZFD-RRM}$ *cmR*]. The genomic DNA of the RIK900 strain was used to transform the strain VHB181, and selected by erythromycin resistance, yielding the desired strain VHB189.

To construct the strain VHB161 [*trpC2 amyE::*$P_{hy-spank}$*-relD264G(1-373)* $^{\Delta CTD}$ *cmR rel::ermR*], plasmid pHT125 was subjected to site-directed mutagenesis using primer VHT69 and VHT190 , according to directions of Phusion Site-Directed Mutagenesis Kit (Thermo Fisher Scientific), yielding plasmid VHP436 (pHT001-*relD264G* $^{\Delta CTD}$) which was used to transform wt168 strain. Selection for chloramphenicol resistance yielded the strain VHB154 [*trpC2 amyE::*$P_{hy-spank}$*-relD264G* $^{\Delta CTD}$ *cmR*]. DNA extracted from strain RIK900 was used to transform the strain VHB154, and selected by erythromycin resistance, yielding the desired strain VHB164. To construct the strain VHB190 [*trpC2 relQ::tet relP::spcR amyE*::$P_{hy-spnak}$-*relD264G* $^{\Delta CTD}$ *cmR rel::ermR* ], plasmid VHP436 was used to transform strain NBS1437 [*trpC2 relQ::tet relP::spcR*], and selected by chloramphenicol resistance, yielding the strain VHB182 [*trpC2 relQ::tet relP::spcR amyE*::P$_{hy\text{-}spnak}$-*relD264G* $^{\Delta CTD}$ *cmR*]. The genomic DNA of the RIK900 strain was used to transform the strain VHB182, and selected by erythromycin resistance, yielding the desired strain VHB190.

To construct the strain VHB231 [*trpC2 relQ::tet relP::spcR amyE*::P$_{hy\text{-}spnak}$-*relH420E* $^{\Delta RRM}$ *cmR rel::ermR*], plasmid VHP435 was subjected to site-directed mutagenesis using primer VHT337 and VHT338 , according to directions of Phusion Site-Directed Mutagenesis Kit (Thermo Fisher Scientific), yielding plasmid VHP591 (pHT001-*relH420E* $^{-\Delta RRM}$ *amp cmR*) which was used to transform strain NBS1437. Selection for chloramphenicol resistance yielded the strain VHB230 [*trpC2 relQ::tet relP::spcR amyE*::P$_{hy\text{-}spnak}$-*relH420E* $^{\Delta RRM}$ *cmR*]. DNA extracted from strain RIK900 (*trpC2 rel::ermR*) was used to transform the strain VHB230, and selected by erythromycin resistance, yielding the desired strain VHB231.

To construct the strain VHB233 [*trpC2 relQ::tet relP::spcR amyE*::P$_{hy\text{-}spnak}$-*relC602AC603A* $^{\Delta RRM}$ *cmR rel::ermR*], VHP435 was subjected to site-directed mutagenesis using primer VHT337 and VHT338 , according to directions of Phusion Site-Directed Mutagenesis Kit (Thermo Fisher Scientific), yielding plasmid VHP592 (pHT001-*relC602AC603A* $^{-\Delta RRM}$ *amp cmR*) which was used to transform strain NBS1437. Selection for chloramphenicol resistance yielded the VHB232 [*trpC2 relQ::tet relP::spcR amyE*::P$_{hy\text{-}spnak}$-*relC602AC603A* $^{\Delta RRM}$ *cmR*]. DNA extracted from strain RIK900 was used to transform the strain VHB232, and selected by erythromycin resistance, yielding the desired strain VHB233.

To construct the strain VHB252 [*trpC2 amyE::P$_{hy\text{-}spank}$-rel(1-155)* $^{\Delta CTD}$ *cmR rel::ermR*], the pHT101 plasmid was subjected to site-directed mutagenesis using primers VHT69 and VHT341, as per manufacturers manual (Phusion Site-Directed Mutagenesis Kit, Thermo Fisher Scientific), yielding the plasmid VHP622 (pHT001-*rel(1-155)* $^{\Delta CTD}$) which was used to transform wt168 strain. Selection for chloramphenicol resistance yielded the strain VHB249 [*trpC2 amyE::P$_{hy\text{-}spank}$-rel(1-155)* $^{\Delta CTD}$ *cmR*]. DNA extracted from strain RIK900 was used to transform the strain VHB249, and selected by erythromycin resistance, yielding the desired strain VHB252.

To construct the strain VHB253 [*trpC2 amyE::P$_{hy\text{-}spank}$-rel(1-196)* $^{\Delta CTD}$ *cmR rel::ermR*], the pHT101 plasmid was subjected to site-directed mutagenesis using primers VHT69 and VHT342, as per manufacturers manual (Phusion Site-Directed Mutagenesis Kit, Thermo Fisher Scientific), yielding the plasmid VHP623 (pHT001-*rel(1-196)* $^{\Delta CTD}$) which was used to transform wt168 strain. Selection for chloramphenicol resistance yielded the strain VHB250 [*trpC2 amyE::P$_{hy\text{-}spank}$-rel(1-196)* $^{\Delta CTD}$ *cmR*]. DNA extracted from strain RIK900 was used to transform the strain VHB250, and selected by erythromycin resistance, yielding the desired strain VHB253.

To construct the strain VHB254 [*trpC2 amyE::P$_{hy\text{-}spank}$-rel(1-336)* $^{\Delta CTD}$ *cmR rel::ermR*], the pHT101 plasmid was subjected to site-directed mutagenesis using primers VHT69 and VHT344, as per manufacturers manual (Phusion Site-Directed Mutagenesis Kit, Thermo Fisher Scientific), yielding the plasmid VHP624 (pHT001-*rel(1-336)* $^{\Delta CTD}$) which was used to transform wt168 strain. Selection for chloramphenicol resistance yielded the strain VHB251 [*trpC2 amyE::P$_{hy\text{-}spank}$-rel(1-336)* $^{\Delta CTD}$ *cmR*]. DNA extracted from strain RIK900 was used to transform the strain VHB251, and selected by erythromycin resistance, yielding the desired strain VHB254.

To construct the strain VHB281 [*trpC2 relQ::tet relP::spcR amyE*::P$_{hy\text{-}spnak}$-*relR619E* $^{\Delta RRM}$ *cmR rel::ermR*], plasmid VHP435 was subjected to site-directed mutagenesis using primer

VHT413 and VHT414 , according to directions of Phusion Site-Directed Mutagenesis Kit (Thermo Fisher Scientific), yielding plasmid VHP626 (pHT001-*relR619E* $^{\Delta RRM}$) which was used to transform strain NBS1437. Selection for chloramphenicol resistance yielded the strain VHB280 [*trpC2 relQ::tet relP::spcR amyE*::$P_{hy\text{-}spnak}$-*relR619E* $^{\Delta RRM}$ *cmR*]. DNA extracted from strain RIK900 (*trpC2 rel::ermR*) was used to transform the strain VHB, and selected by erythromycin resistance, yielding the desired strain VHB281.

To construct the strain VHB282 [*trpC2 amyE::*$P_{hy\text{-}spank}$-*relR619E cmR rel::ermR*], plasmid pHT101 was subjected to site-directed mutagenesis using primer VHT413 and VHT414 , according to directions of Phusion Site-Directed Mutagenesis Kit (Thermo Fisher Scientific), yielding the plasmid VHP625 (pHT001-*relR619E*) which was used to transform wt168 strain. Selection for chloramphenicol resistance yielded the strain VHB273 [*trpC2 amyE::*$P_{hy\text{-}spank}$-*relR619E cmR*]. DNA extracted from strain RIK900 was used to transform the strain VHB273, and selected for erythromycin resistance, yielding the desired strain VHB282.

Strain VHB455 [*trpC2 relN685A-spcR*] was constructed using the site-directed mutagenesis technique. To exchange the AAT codon (coding for Asparagine) at position 685 for GCG (coding for Alanine), two ≈2,500 nucleotide-long overlapping DNA fragments were amplified by PCR using genomic DNA of strain NBS1729 strain as a template. The first fragment used as the DNA template the region located upstream of the *rel* ORF, spanning to codon 685 of the *rel* ORF (primers: VHT29 and VHT83). The second used as a template the region starting with the codon 685 of *rel* ORF, spanning downstream of the *rel* ORF to include the *spcR* resistance marker (primers: VHT30 and VHT82). The two PCR products described above were used simultaneously as the template for PCR amplification using primers VHT29 and VHT30. The resulting long fragment was used to transform wild-type strain 168 (*trpC2*). The transformants were selected by spectinomycin resistance yielding the strain VHB455. The desired mutation was confirmed by sequencing.

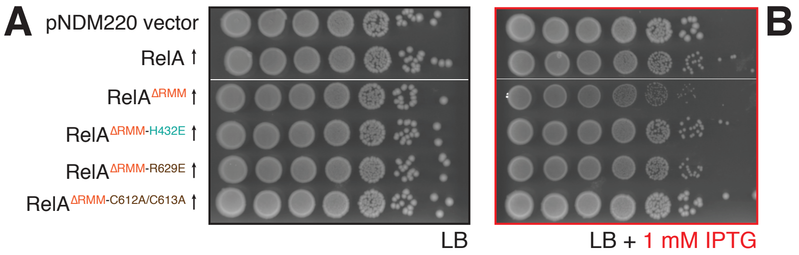


**Supplementary Figure S1. The toxicity of ΔRRM *E. coli* RelA expressed in the Δ*relA* background is mitigated by mutations compromising interactions with tRNA and the ribosome.** *E. coli* BW25113 Δ*relA* cells were transformed with low copy IPTG-inducible vector pNDM220 (vector) or pNDM220-based constructs expressing either wild-type and mutant versions of *E. coli* RelA as indicated on the figure. Ten-fold serial dilutions of overnight LB cultures were made and spotted onto LB agar (LB) supplemented with 30 µg/mL ampicillin, without (**A**, left panel) and with 1 mM IPTG (**B**, right panel). The plates were incubated at 37 °C and scored after 18 hours. Growth assays presented on panels **A** and **B** originate from the same individual plates after cropping and splicing together.

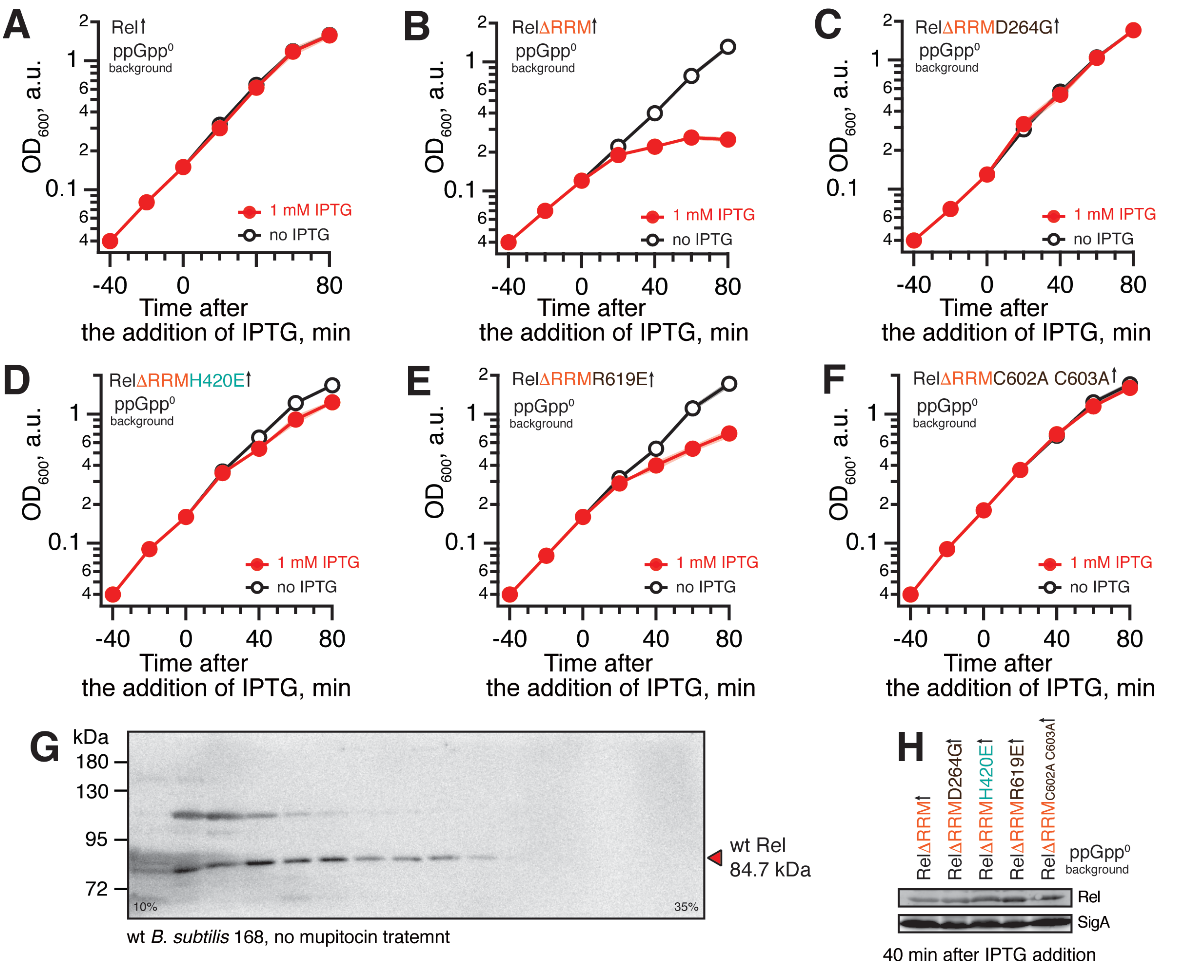


**Supplementary Figure S2. The toxicity of *B. subtilis* ΔRRM Rel is mediated by (p)ppGpp synthesis and is countered by mutations compromising the interaction with starved ribosomes.** Full-length (**A**: VHB155) or C-terminally ΔRRM truncated Rel variants (**B**: VHB184; **C**: VHB188; **D**: VHB231; **E**: VHB281; **F**: VHB233) were expressed in ppGpp$^0$ *B. subtilis* growing in liquid LB medium at 37 °C. The error bars represent standard deviation of the mean (n = 3) and are shown as shading since they are generally smaller than the markers. (**G**) Uncut version of the immunoblot analysis presented on **Figure 4A**, top line. (**H**) anti-Rel and anti-SigA immunoblot analyses of samples collected 40 minutes upon induction of expression Rel$^{\Delta RRM}$ or D264G, H420E, R619E and C602A C603A Rel$^{\Delta RRM}$ variants by IPTG.

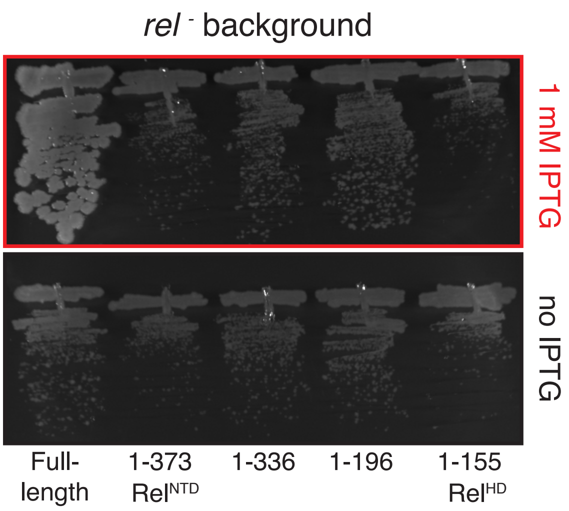


**Supplementary Figure S3. None of the tested truncated variants of Rel$^{NTD}$ rescue the growth effect of Δ*rel B. subtilis.*** Either full-length or N-terminally truncated Rel variants were expressed in Δ*rel B. subtilis* (VHB155, VHB161, VHB252-254) growing on solid LB medium at 37 °C and plates were scored after 18 hour incubation.

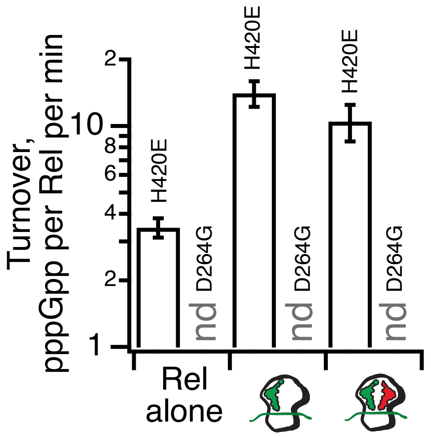


**Supplementary Figure S4. *B. subtilis* D264G Rel is catalytically inactive and *B. subtilis* H420E Rel is not activated by A-site deacylated tRNA.** $^{3}$H-pppGpp synthesis activity of 140 nM of Rel H420E and Rel D264G assayed in the presence of 1 mM ATP nd 0.3 mM of either $^{3}$H-labelled GTP. As indicated on the figure, the reaction mixtures were supplemented with combinations of 0.5 µM IC (MV) and native *E. coli* $tRNA^{Val}$ (2 µM: A-site) as well as 100 µM pppGpp. All experiments were performed in HEPES:Polymix buffer, pH 7.5 at 37 °C in the presence of 5 mM $Mg^{2+}$. Error bars represent SDs of the turnover estimates by linear regression using four data points.

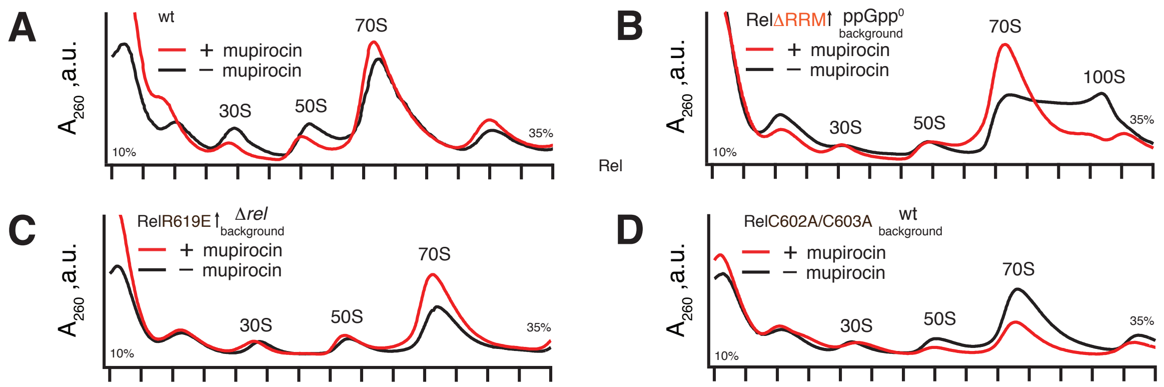


**Supplementary Figure S5.** Polysome profiles of Rel variants expressed either ectopically under the control of IPTG-inducible $P_{hy\text{-}spnak}$ promotor [ΔRRM Rel in ppGpp$^0$ *B. subtilis* (VHB184), R619E Rel in Δ*rel B. subtilis* (VHB282)] or from the native chromosomal locus [C602A C603A mutant (VHB144)]. Expression of Rel$^{\Delta RRM}$ was induced by 1 mM IPTG for 10 minutes followed by a 10 minute challenge with 700 nM mupirocin. To drive the expression of RelR619E Rel the strain as grown in LB supplemented with 1 mM IPTG. In the case of R619E and C602AC603A Rel the culture was treated with mupirocin for 20 minutes.

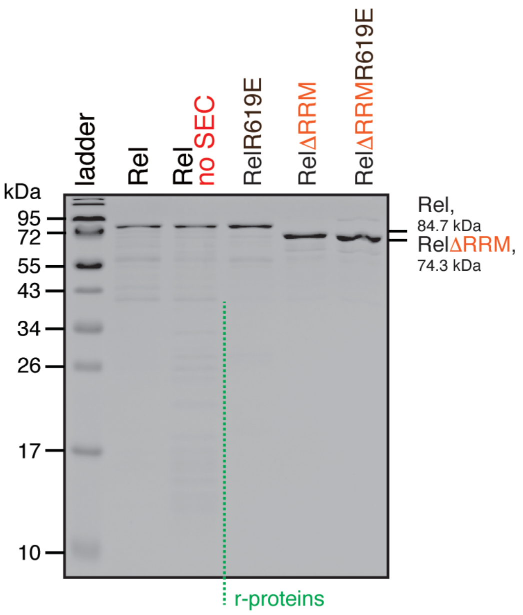


**Supplementary Figure S6. SDS-PAGE analysis of wild-type Rel as well as ΔRRM, R619E and ΔRRM R619E mutant variants.** The full-length Rel protein was purified either as described on **Figure 5** or with the SEC step omitted (no SEC).

**Supplementary Table 1.** ***E. coli*** **strains and plasmids used in this study.**

| **Strain / plasmid** | **Description** | **Reference** |
|---|---|---|
| BL21 DE | BF⁻ *ompT gal dcm lon* $hsdS_B(r_B^- m_B^-)$ λ(DE3 [*lacI lacUV5-T7p07 ind1 sam7 nin5*]) [$malB^+$]$_{K\text{-}12}$($\lambda^S$) | laboratory collection |
| BW25113 | wild type *E. coli* K-12 | laboratory collection |
| BW25113 **Δ*relA*** | Δ*relA E. coli* BW25113 | (Varik et al., 2016) |
| pNDM220 | mini-R1 $lacI^q$ $P_{A1/O4/O3}$ $amp^R$ | (Gotfredsen and Gerdes, 1998) |
| pNDM220::*relA* | pNDM220::*relA* full-length (residues 1-744) | (Turnbull et al., 2019) |
| pNDM220::$relA^{\Delta RRM}$ | pNDM220::*relA* ΔRRM domain (residues 1-657) | (Turnbull et al., 2019) |
| pNDM220::$relA^{\Delta RRM\text{-}H432E}$ | pNDM220::*relA* ΔRRM, H432E mutant | this work |
| pNDM220::$relA^{\Delta RRM\text{-}R629E}$ | pNDM220::*relA* ΔRRM, R629E mutant | this work |
| pNDM220::$relA^{\Delta RRM\text{-}C612A/C613A}$ | pNDM220::*relA* ΔRRM, C612A/C613A mutant | this work |
| pMG25 | pUC $lacI^q$ $P_{A1/O4/O3}$ $amp^R$ | (Turnbull et al., 2019) |
| pMG25::*relA* | pMG25::*relA* full-length (residues 1-744) | (Turnbull et al., 2019) |
| pMG25::$relA^{\Delta RRM}$ | pMG25::*relA* ΔRMM domain (residues 1-657) | this work |
| pMG25::*spoT* | pMG25::*spoT* full-length (residues 1-702) | this work |
| pMG25::$spoT^{\Delta RRM}$ | pMG25:: *spoT* ΔRMM domain (residues 1-617) | this work |
| pET24::*relA* | N-terminal $His_{10}$-SUMO::*relA* | (Turnbull et al., 2019) |
| pET24::$relA^{H432E}$ | N-terminal $His_{10}$-SUMO::$relA^{H432E}$ | this work |

**Supplementary Table 2. *B. subtilis* strains used in this study.**

| Strain | Description | Reference |
|---|---|---|
| wild type 168 | *trpC2* | laboratory stock |
| VHB47 | *trpC2 rplK::cmR* | this work |
| VHB49 | *trpC2 relQ::tet relP::spcR rplK::cmR* | this work |
| VHB60 | *trpC2ΔrelQ relP::cmR relH420E-spcR* | this work |
| VHB62 | *trpC2 ΔrelQ relP::cmR relC602AC603A-spcR* | this work |
| VHB63 | *trpC2 ΔrelQ relP::cmR rel:::spcR* | this work |
| VHB68 | *trpC2 relH420E-spcR* | this work |
| VHB144 | *trpC2 relC602AC603A-spcR* | this work |
| VHB145 | *trpC2 amyE*::$P_{hy\text{-}spnak}$*-rel cmR* | this work |
| VHB146 | *trpC2 amyE*::$P_{hy\text{-}spnak}$*-rel*$^{\Delta RRM}$ *cmR* | this work |
| VHB147 | *trpC2 amyE*::$P_{hy\text{-}spnak}$*-rel*$^{\Delta ZFD\text{-}RRM}$ *cmR* | this work |
| VHB148 | *trpC2 amyE*::$P_{hy\text{-}spnak}$*-rel(1-373)*$^{NTD}$ *cmR* | this work |
| VHB151 | *trpC2 amyE*::$P_{hy\text{-}spnak}$*-relD264G cmR* | this work |
| VHB152 | *trpC2 amyE*::$P_{hy\text{-}spnak}$*-relD264G*$^{\Delta RRM}$ *cmR* | this work |
| VHB153 | *trpC2 amyE*::$P_{hy\text{-}spnak}$*-relD264G*$^{\Delta ZFD\text{-}RRM}$ *cmR* | this work |
| VHB154 | *trpC2 amyE*::$P_{hy\text{-}spnak}$*-relD264G(1-373)*$^{NTD}$ *cmR* | this work |
| VHB155 | *trpC2 amyE*::$P_{hy\text{-}spnak}$*-rel cmR rel::ermR* | this work |
| VHB156 | *trpC2 amyE*::$P_{hy\text{-}spnak}$*-relD264G cmR rel::ermR* | this work |
| VHB159 | *trpC2 amyE*::$P_{hy\text{-}spnak}$*-rel*$^{\Delta RRM}$ *cmR rel::ermR* | this work |
| VHB160 | *trpC2 amyE*::$P_{hy\text{-}spnak}$*-rel*$^{\Delta ZFD\text{-}RRM}$ *cmR rel::ermR* | this work |
| VHB161 | *trpC2 amyE*::$P_{hy\text{-}spnak}$*-rel*$^{NTD}$ *cmR rel::ermR* | this work |
| VHB162 | *trpC2 amyE*::$P_{hy\text{-}spnak}$*-relD264G* $^{\Delta RRM}$ *cmR rel::ermR* | this work |
| VHB163 | *trpC2 amyE*::$P_{hy\text{-}spnak}$*-relD264G*$^{\Delta ZFD\text{-}RRM}$ *cmR rel::ermR* | this work |
| VHB164 | *trpC2 amyE*::$P_{hy\text{-}spnak}$*-relD264G(1-373)*$^{NTD}$ *cmR rel::ermR* | this work |
| VHB175 | *trpC2 relQ::tet relP::spcR amyE*::$P_{hy\text{-}spnak}$*-rel cmR* | this work |
| VHB176 | *trpC2 relQ::tet relP::spcR amyE*::$P_{hy\text{-}spnak}$*-rel*$^{\Delta RRM}$ *cmR* | this work |
| VHB177 | *trpC2 relQ::tet relP::spcR amyE*::$P_{hy\text{-}spnak}$*-rel*$^{\Delta ZFD\text{-}RRM}$ *cmR* | this work |
| VHB178 | *trpC2 relQ::tet relP::spcR amyE*::$P_{hy\text{-}spnak}$*-rel(1-373)*$^{NTD}$ *cmR* | this work |
| VHB179 | *trpC2 relQ::tet relP::spcR amyE*::$P_{hy\text{-}spnak}$*-relD264G cmR* | this work |
| VHB180 | *trpC2 relQ::tet relP::spcR amyE*::$P_{hy\text{-}spnak}$*-relD264G*$^{\Delta RRM}$ *cmR* | this work |
| VHB181 | *trpC2 relQ::tet relP::spcR amyE*::$P_{hy\text{-}spnak}$*-relD264G*$^{\Delta ZFD\text{-}RRM}$ *cmR* | this work |
| VHB182 | *trpC2 relQ::tet relP::spcR amyE*::$P_{hy\text{-}spnak}$*-relD264G(1-373)*$^{NTD}$ *cmR* | this work |
| VHB183 | *trpC2 relQ::tet relP::spcR amyE*::$P_{hy\text{-}spnak}$*-rel cmR rel::ermR* | this work |
| VHB184 | *trpC2 relQ::tet relP::spcR amyE*::$P_{hy\text{-}spnak}$*-rel*$^{\Delta RRM}$ *cmR rel::ermR* | this work |
| VHB185 | *trpC2 relQ::tet relP::spcR amyE*::$P_{hy\text{-}spnak}$*-rel*$^{\Delta ZFD\text{-}RRM}$ | this work |

| | | |
|---|---|---|
| | *cmR rel::ermR* | |
| VHB186 | *trpC2 relQ::tet relP::spcR amyE*::$P_{hy\text{-}spnak}$-*rel(1-373)*$^{NTD}$ *cmR rel::ermR* | this work |
| VHB187 | *trpC2 relQ::tet relP::spcR amyE*::$P_{hy\text{-}spnak}$-*relD264G cmR rel::ermR* | this work |
| VHB188 | *trpC2 relQ::tet relP::spcR amyE*::$P_{hy\text{-}spnak}$-*relD264G*$^{\Delta RRM}$ *cmR rel::ermR* | this work |
| VHB189 | *trpC2 relQ::tet relP::spcR amyE*::$P_{hy\text{-}spnak}$-*relD264G*$^{\Delta ZFD\text{-}RRM}$ *cmR rel::ermR* | this work |
| VHB190 | *trpC2 relQ::tet relP::spcR amyE*::$P_{hy\text{-}spnak}$-*relD264G(1-373)*$^{NTD}$ *cmR rel::ermR* | this work |
| VHB230 | *trpC2 relQ::tet relP::spcR amyE*::$P_{hy\text{-}spnak}$-*relH420E*$^{\Delta RRM}$ *cmR* | this work |
| VHB231 | *trpC2 relQ::tet relP::spcR amyE*::$P_{hy\text{-}spnak}$-*relH420E*$^{\Delta RRM}$ *cmR rel::ermR* | this work |
| VHB232 | *trpC2 relQ::tet relP::spcR amyE*::$P_{hy\text{-}spnak}$-*relC602AC603A*$^{\Delta RRM}$ *cmR* | this work |
| VHB233 | *trpC2 relQ::tet relP::spcR amyE*::$P_{hy\text{-}spnak}$-*relC602AC603A*$^{\Delta RRM}$ *cmR rel::ermR* | this work |
| VHB249 | *trpC2 amyE*::$P_{hy\text{-}spnak}$-*rel(1-155)*$^{NTD}$ *cmR* | this work |
| VHB250 | *trpC2 amyE*::$P_{hy\text{-}spnak}$-*rel(1-196)*$^{NTD}$ *cmR* | this work |
| VHB251 | *trpC2 amyE*::$P_{hy\text{-}spnak}$-*rel(1-336)*$^{NTD}$ *cmR* | this work |
| VHB252 | *trpC2 amyE*::$P_{hy\text{-}spnak}$-*rel(1-155)*$^{NTD}$ *cmR rel::ermR* | this work |
| VHB253 | *trpC2 amyE*::$P_{hy\text{-}spnak}$-*rel(1-196)*$^{NTD}$ *cmR rel::ermR* | this work |
| VHB254 | *trpC2 amyE*::$P_{hy\text{-}spnak}$-*rel(1-336)*$^{NTD}$ *cmR rel::ermR* | this work |
| VHB273 | *trpC2 amyE*::$P_{hy\text{-}spnak}$-*relR619E cmR* | this work |
| VHB280 | *trpC2 relQ::tet relP::spcR trpC2 amyE*::$P_{hy\text{-}spnak}$-*relR619E*$^{\Delta RRM}$ *cmR* | this work |
| VHB281 | *trpC2 relQ::tet relP::spcR trpC2 amyE*::$P_{hy\text{-}spnak}$-*relR619E*$^{\Delta RRM}$ *cmR rel::ermR* | this work |
| VHB282 | *trpC2 amyE*::$P_{hy\text{-}spnak}$-*relR619E cmR rel::ermR* | this work |
| VHB455 | *trpC2 relN685A-spcR* | this work |
| RIK900 | *trpC2 rel::ermR* | (Nanamiya et al., 2008) |
| RIK908 | *trpC2 relP::spcR* | (Nanamiya et al., 2008) |
| RIK1000 | *trpC2 ΔrelQ* | (Nanamiya et al., 2008) |
| RIK1004 | *trpC2 relQ::cmR* | (Tagami et al., 2012) |
| RIK2508 | *trpC2 Δhpf* | (Akanuma et al., 2016) |
| NBS1393 | *trpC2 relQ::tet* | this work |
| NBS1437 | *trpC2 relQ::tet relP::spcR* | this work |
| NBS1729 | *trpC2 rel-spcR* | this work |
| NHT436 | *trpC2 ΔrelQ relP::cmR* | this work |
| NHT437 | *trpC2 rel::spcR* | this work |

**Supplementary Table 3. *B. subtilis* plasmids used in this study.**

| Plasmid | Description | Reference |
|---|---|---|
| pHT101 | pHT001-*rel amp cmR* | this work |
| pHT125 | pHT001-*relD264G amp cmR* | this work |
| VHP186 | pET24d-*His$_{10}$-SUMO-rel kmR* | this work |
| VHP187 | pET24d-*His$_{10}$-SUMO-relD264G kmR* | this work |
| VHP230 | pET24d-*His$_{10}$-SUMO-relH420E kmR* | this work |
| VHP231 | pET24d-*His$_{10}$-SUMO-rel$^{\Delta RRM}$ kmR* | this work |
| VHP433 | pHT001-*rel(1-373)$^{NTD}$ amp cmR* | this work |
| VHP434 | pHT001-*rel$^{\Delta ZFD\text{-}RRM}$ amp cmR* | this work |
| VHP435 | pHT001-*rel$^{\Delta RRM}$ amp cmR* | this work |
| VHP436 | pHT001-*relD264G(1-373) $^{NTD}$ amp cmR* | this work |
| VHP437 | pHT001-*relD264G$^{\Delta ZFD\text{-}RRM}$ amp cmR* | this work |
| VHP438 | pHT001-*relD264G$^{\Delta RRM}$ amp cmR* | this work |
| VHP591 | pHT001-*relH420E$^{\Delta RRM}$ amp cmR* | this work |
| VHP592 | pHT001-*relC602AC603A$^{\Delta RRM}$ amp cmR* | this work |
| VHP622 | pHT001-*rel(1-155)$^{NTD}$ amp cmR* | this work |
| VHP623 | pHT001-*rel(1-196)$^{NTD}$ amp cmR* | this work |
| VHP624 | pHT001-*rel(1-336)$^{NTD}$ amp cmR* | this work |
| VHP625 | pHT001-*relR619E amp cmR* | this work |
| VHP626 | pHT001-*relR619E$^{\Delta RRM}$ amp cmR* | this work |
| VHP627 | pET24d-*His$_{10}$-SUMO-relR619E kmR* | this work |
| VHP630 | pET24d-*His$_{10}$-SUMO-relR619E$^{\Delta RRM}$ kmR* | this work |

**Supplementary Table 4. Primers used in this work.**

| Primer | Sequence, 5'-3' |
|---|---|
| VHT1 | ATTAAGTGATTTAGAGAGAAAGTACTCGTCTTATATCTCG |
| VHT2 | AAGAGCAAGGTTGCGAACTGGA |
| VHT5 | ATCTAAAGTGTTAGCTGGGTTAGCTTTTCCAGCAG |
| VHT6 | CTAACCCAGCTAACACTTTAGATAAAAATTTAGGAGGC |
| VHT9 | GCTTTTATAACGCAGTATGGGTATTGTAATCGAGGATTAA |
| VHT10 | CCATACTGCGTTATAAAAGCCAGTCATTAGGCC |
| VHT29 | AAGATCAAGCAAAGGCGTTTGACCTCGAT |
| VHT30 | GTGCGTCAACGACGGGCACAGTC |
| VHT31 | TTACCGGATTGAGTCTGAAATCGGCAATAAAACAATCGGT |
| VHT32 | CGATTTCAGACTCAATCCGGTAAGAAAAGTCAATCGGAAC |
| VHT39 | GTTTATCAAAAGCCGCCAATCCTGTGCCAGGTGATGATATTGTC |
| VHT40 | GGCACAGGATTGGCGGCTTTTGATAAACGGACAAGGAGGTTGTCAAT |
| VHT68 | CCACTCTACCGGGATCAGCCGCT |
| VHT69 | TAAGCATGCAAGCTAATTCGGTGGAAACG |
| VHT82 | GAAAACCGCGATTTCATCTGTCTCTGGCAAATCGGATCG |
| VHT83 | CAGATGAAATCGCGGTTTTCGTTTCATTCACTGCCTGGAG |
| VHT123 | CATTATCGCTCTCTCCTTCGTCGACTAAGCTAATTG |
| VHT125 | TAAGCATGCAAGCTAATTCGGTGGAAACGAGG |
| VHT131 | GGCGGCCATCATCATCATCATCACGCCAA |
| VHT169 | CGACGAAGGAGAGAGCGATAATGGCGAACGAACAAGTATTG |
| VHT170 | TGATGATGATGATGGCCGCCGTTCATGACGCGGCGCA |
| VHT189 | CCGTTTTCTTCCTTGCGGGTATGGCT |
| VHT190 | ATTTTGAAATTCTAAAATTTCACGGAACCAAGAAAGCTTTTTCTCAAAG |
| VHT337 | CTCAATCCGGTAAGAAAAGTCAATCGGAACAGAACCGGA |
| VHT338 | TCTGAAATCGGCAATAAAACAATCGGTGCCAAAGT |
| VHT339 | GCCAATCCTGTGCCAGGTGATGATATTGTCGGCT |

| | |
|---|---|
| VHT340 | GGCTTTTGATAAACGGACAAGGAGGTTGTCAATGCCCT |
| VHT341 | CAGATGTTTCAGTGTCCGCATATTGTGAAGACGAT |
| VHT342 | GTTCAAATAACGGAGCGCCGTATCTTCCAATTCCCA |
| VHT344 | CCGCTATTTCATGCATTTCAAAGGTGCGGA |
| VHT383 | CGACGAAGGAGAGAGCGATAATGGTTGCGGTAAGAAGTGCA |
| VHT384 | CACCGAATTAGCTTGCATGCTCAACTCCCGTGCAACCGA |
| VHT385 | CACCGAATTAGCTTGCATGCTCACCATACCGCGTCAACAAT |
| VHT386 | CGACGAAGGAGAGAGCGATAATGCAAGTTATTGACTCTAAACT |
| VHT387 | CACCGAATTAGCTTGCATGCTTAGCTGACCTCTTCATTATCATCCCA |
| VHT413 | GAAGGGGTTTCGGTCCATCGCGAAGACTGT |
| VHT414 | GCCTTTTGTGATAAAGCCGACAATATCATCACCTG |
| NOP1 | GCGGCGTGAAGAAAATGTCG |
| NOP2 | CACCTCGTTGTTAGTTCATGACGCGGCGCA |
| NOP3 | CGTCATGAACTAACAACGAGGTGAAATCATGAG |
| NOP4 | TCTAACCCCTTTACTAGGCCTAATTGAGAGAAG |
| NOP5 | CAATTAGGCCTAGTAAAGGGGTTAGAAAAGAGATTAG |
| NOP6 | CAGTCCTGCCAAAGATCAGGAAC |
| NOP7 | ATTGAACTGGCAGGCCT |
| NOP8 | TACCTCTGAAAGCTGCTGG |
| NOP9 | GGGATGTATGTAGCGGTTATGAAGTGAAATTGA |
| NOP10 | TCCCCTTTACTTACTAGAAATCCCTTTGAGAATG |
| NOP11 | AGGGATTTCTAGTAAGTAAAGGGGAAGAAGAGCATG |
| NOP12 | TTTCGGATGCAACTTGGC |
| NOP13 | CTCGTTGTTATGCTCGGCAGTCAATACTTGTTCGT |
| NOP14 | CTGCCGAGCATAACAACGAGGTGAAATCATGAG |
| NOP15 | AGAATAGATATTACTAGGCCTAATTGAGAGAAG |
| NOP16 | GGCCTAGTAATATCTATTCTGTGCGCCGCGTCA |
| NSP1 | ACATGCATGCTTAGTTCATGACGCGGC |

| | |
|---|---|
| NSP2 | ACGCGTCGACCTTGATAGTAAACTACAATACAGTTTACA |
| NHP1 | GGCTTGTTGGCTGTCCGTATTCTTGTGAATAGC |
| NHP2 | GTAAATTTCATTGAATTGCTTATTTGCAGCACCATTTTTCG |
| pQErelA_F | GCCCCATGGATGGCGAACGAACAAGTATTG |
| pQErelA_R | GTCCCATGGGTTCATGACGCGGCGCACAG |
| MR1 | CCCCCGGATCCTGGTCCCTAAAGGAGAGG |
| MR2 | CCCCCCTCGAGTTACCATACCGCGTCAACAATGC |
| MR3 | GGACTTCGCTTACCACATCGAGAGTGATGTCGGACACCGCTG |
| MR4 | CAGCGGTGTCCGACATCACTCTCGATGTGGTAAGCGAAGTCC |
| MR5 | GCTTCATTACCCAGGGGGAGGGTATTTCAGTACACCG |
| MR6 | CGGTGTACTGAAATACCCTCCCCCTGGGTAATGAAGC |
| MR7 | GCACCACATCGCGCGCGCCGCCCAGCCGATTCCTGGAG |
| MR8 | CTCCAGGAATCGGCTGGGCGGCGCGCGCGATGTGGTGC |
